# Identification of novel rare sequence variation underlying heritable pulmonary arterial hypertension

**DOI:** 10.1101/185272

**Authors:** Stefan Gräf, Matthias Haimel, Marta Bleda, Charaka Hadinnapola, Laura Southgate, Wei Li, Joshua Hodgson, Bin Liu, Richard M. Salmon, Mark Southwood, Rajiv D. Machado, Jennifer M. Martin, Carmen M. Treacy, Katherine Yates, Louise C. Daugherty, Olga Shamardina, Deborah Whitehorn, Simon Holden, Micheala Aldred, Harm J. Bogaard, Colin Church, Gerry Coghlan, Robin Condliffe, Paul A. Corris, Cesare Danesino, Mélanie Eyries, Henning Gall, Stefano Ghio, Hossein-Ardeschir Ghofrani, J. Simon R. Gibbs, Barbara Girerd, Arjan C. Houweling, Luke Howard, Marc Humbert, David G. Kiely, Gabor Kovacs, Robert V. MacKenzie Ross, Shahin Moledina, David Montani, Michael Newnham, Andrea Olschewski, Horst Olschewski, Andrew J. Peacock, Joanna Pepke-Zaba, Inga Prokopenko, Christopher J. Rhodes, Laura Scelsi, Werner Seeger, Florent Soubrier, Dan F. Stein, Jay Suntharalingam, Emilia Swietlik, Mark R. Toshner, Anton Vonk Noordegraaf, David A. van Heel, Quinten Waisfisz, John Wharton, Stephen J. Wort, Willem H. Ouwehand, Nicole Soranzo, Allan Lawrie, Paul D. Upton, Martin R. Wilkins, Richard C. Trembath, Nicholas W. Morrell

## Abstract

Pulmonary arterial hypertension (PAH) is a rare disorder with a poor prognosis. Deleterious variation within components of the transforming growth factor-β pathway, particularly the bone morphogenetic protein type 2 receptor (*BMPR2*), underlie most heritable forms of PAH. Since the missing heritability likely involves genetic variation confined to small numbers of cases, we performed whole genome sequencing in 1038 PAH index cases and 6385 PAH-negative control subjects. Case-control analyses revealed significant overrepresentation of rare variants in novel genes, namely *ATP13A3, AQP1* and *SOX17*, and provided independent validation of a critical role for *GDF2* in PAH. We provide evidence for familial segregation of mutations in *SOX17* and *AQP1* with PAH. Mutations in *GDF2*, encoding a BMPR2 ligand, led to reduced secretion from transfected cells. In addition, we identified pathogenic mutations in the majority of previously reported PAH genes, and provide evidence for further putative genes. Taken together these findings provide new insights into the molecular basis of PAH and indicate unexplored pathways for therapeutic intervention.

## Introduction

Idiopathic and heritable pulmonary arterial hypertension (PAH) are rare disorders characterised by occlusion of arterioles in the lung^1^, leading to marked increases in pulmonary vascular resistance^2^. Life expectancy from diagnosis averages 3-5 years^3^, with death ensuing from failure of the right ventricle.

Mutations in the gene encoding the bone morphogenetic protein type 2 receptor (*BMPR2*), a receptor for the transforming growth factor-beta (TGF-β) superfamily^4,5^ account for over 80% of families with PAH, and approximately 20% of sporadic cases^6^. Mutations have been identified in genes encoding other components of the TGF-β/bone morphogenetic protein (BMP) signalling pathways, including *ACVRL1*^7^ and *ENG*^8^. On endothelial cells, BMPR2 and ACVRL1 form a signaling complex, utilizing ENG as a co-receptor. Case reports of rare sequence variation in the BMP signalling intermediaries, *SMAD1*, *SMAD4* and *SMAD9*^9,10^, provide compelling evidence for a central role of dysregulated BMP signalling in PAH pathogenesis.

Analysis of coding variation in *BMPR2*-negative kindreds revealed heterozygous mutations in genes not directly impacting on the TGF-β/BMP pathway, including *CAV1*^11^, and the potassium channel, *KCNK3*^12^. Deletions and loss of function mutations in *TBX4*, an essential regulator of embryonic development, were identified in childhood onset PAH^13^. A clinically and pathologically distinct form of PAH, known as pulmonary veno-occlusive disease or pulmonary capillary haemangiomatosis (PVOD/PCH), was shown recently to be caused by biallelic recessive mutations in *EIF2AK4*^14, 15^, a kinase in the integrated stress response.

The purpose of the present study was to identify additional rare sequence variation contributing to the genetic architecture of PAH, and to assess the relative contribution of rare variants in genes implicated in prior studies. A major finding is that rare likely causal heterozygous variants in several previously unidentified genes (*ATP13A3, AQP1* and *SOX17*) were significantly overrepresented in the PAH cohort, and we provide independent validation for *GDF2* as a causal gene.

## Results

### Description of the PAH cohort

In total, 1048 PAH cases (1038 index cases and 10 related affected individuals) were recruited for WGS. Of these, 908 (86.7%) were diagnosed with idiopathic PAH, 58 (5.5%) gave a family history of PAH and 60 (5.7%) gave a history of drug exposure associated with PAH^16^. Twenty two cases (2.1%) held a clinical diagnosis of PVOD/PCH (Figure 1a). Demographic and clinical characteristics of the PAH cohort are provided in Supplementary Table 1. An additional UK family was recruited separately for novel gene identification studies. Briefly, the proband was diagnosed at 12 years with a persistent ductus arteriosus and elevated pulmonary arterial pressure. Explant lung histology following heart-lung transplantation revealed the presence of plexiform lesions. Two of the proband’s offspring were also diagnosed with childhood-onset PAH, one of which had an atrial septal defect. The proband’s parents, siblings and a third child showed no evidence of cardiovascular disease.

**Figure 1.**
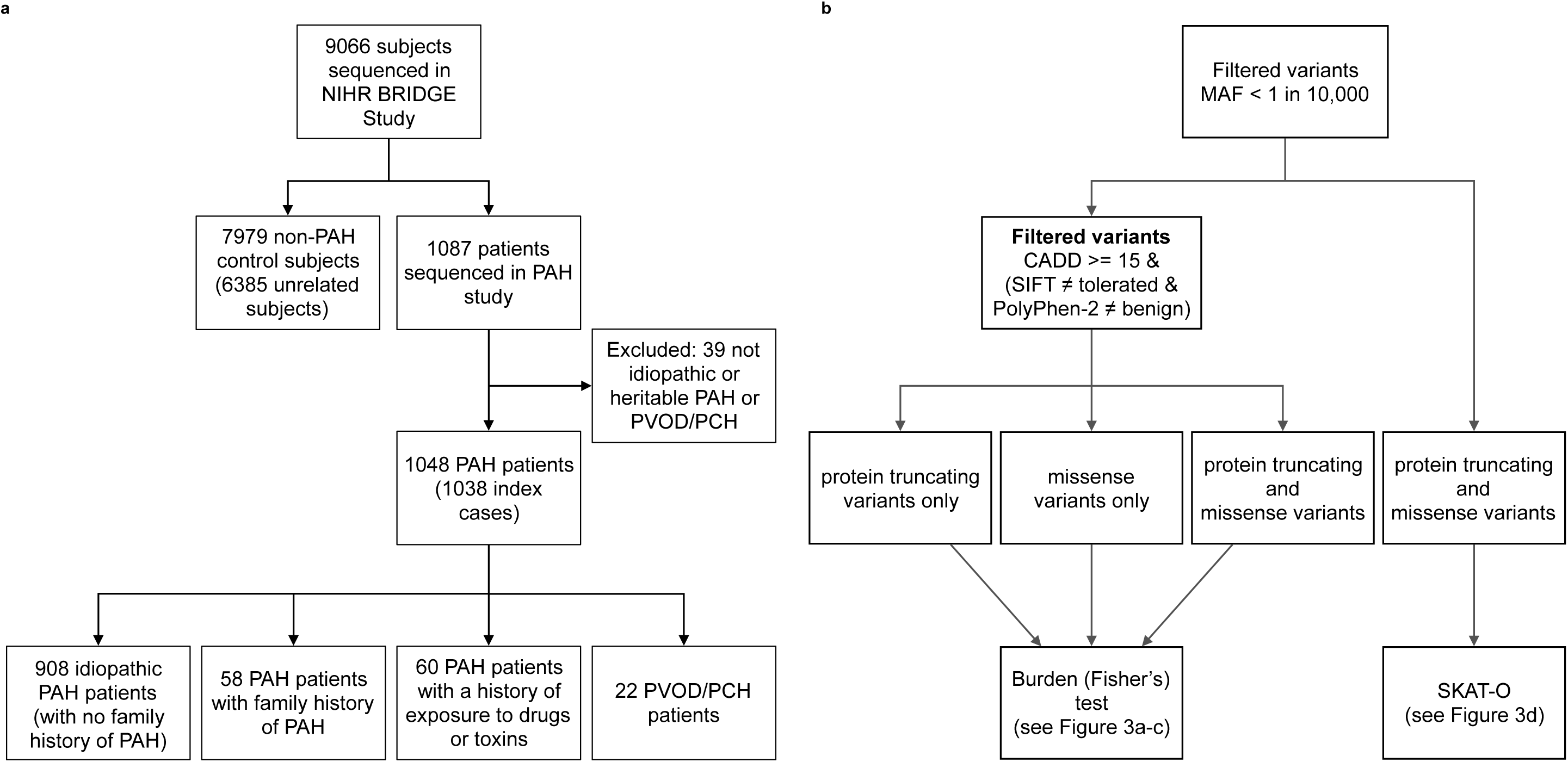
Flow diagrams illustrating **(a)** the composition of the NIHR BioResource – Rare Diseases PAH study and **(b)** the analysis strategy to identify novel PAH disease genes. **(a)** The study comprised 1048 adult cases (aged 16 or over) attending specialist pulmonary hypertension centres from the UK (n=731), and additional cases from France (n=142), The Netherlands (n=45), Germany (n=82) and Italy (n=48). **(b)** A series of case-control comparisons including and excluding cases with variants in previously reported disease genes were undertaken using complementary filtering strategies.

### Pathogenic variants in previously reported PAH disease genes

Our filtering strategy detected rare deleterious variation in previously reported PAH genes in 19.9% of the PAH cohort. For *BMPR2*, rare heterozygous mutations were identified in 160 of 1048 cases (15.3%). The frequency of *BMPR2* mutations in familial PAH was 75.9%, in sporadic cases 12.2%, and 8.3% in anorexigen-exposed PAH cases. Forty-eight percent of *BMPR2* mutations were reported previously^17^, and the remainder were newly identified in this study. Fourteen percent of *BMPR2* mutations resulted in the deletion of larger protein-coding regions ranging from 5 kb to 3.8 Mb in size. Supplementary Table 2 provides the breakdown of *BMPR2* SNVs and indels, and the larger deletions are shown in Figure 2a-c with a detailed summary in Supplementary Table 3.

**Figure 2.**
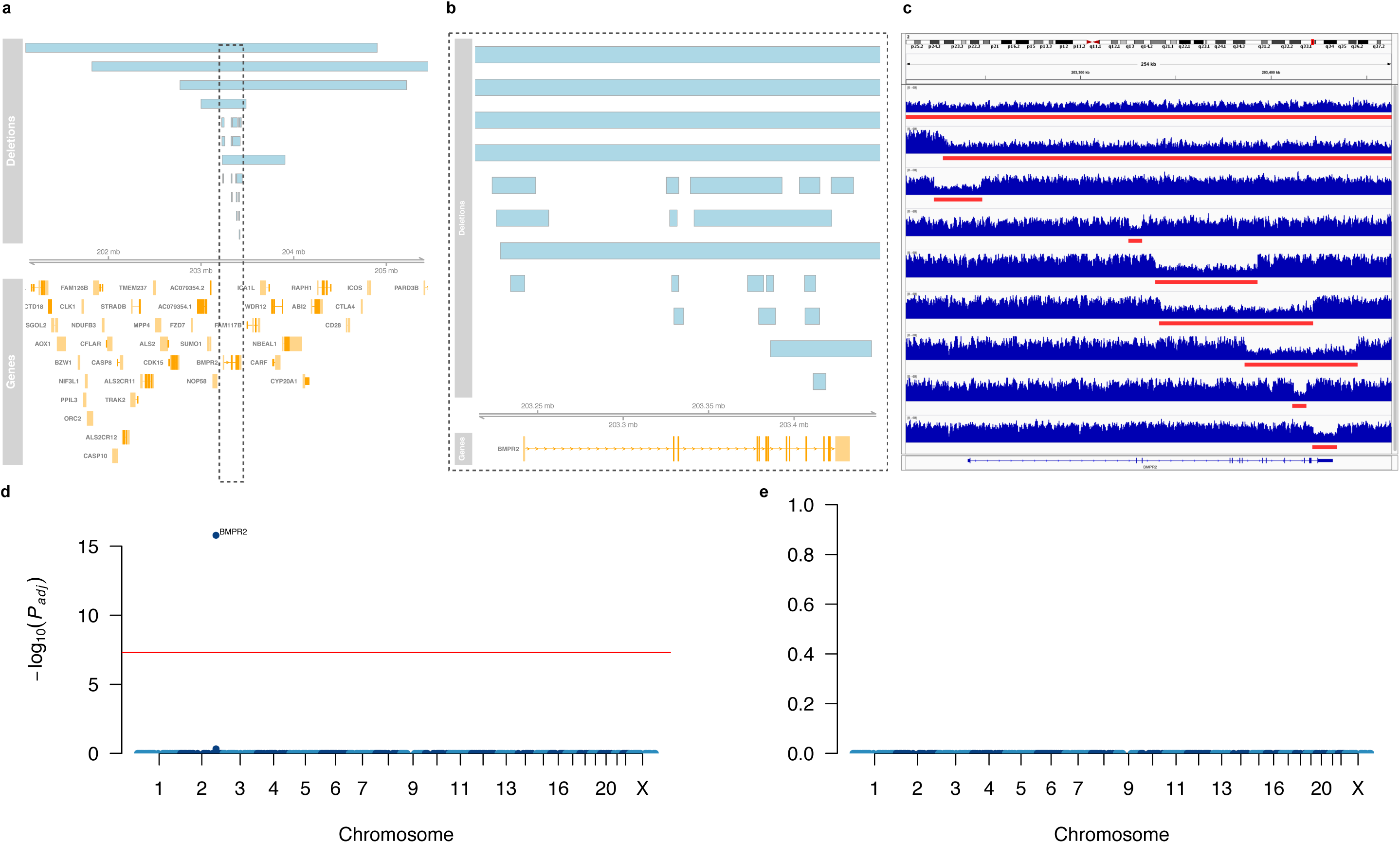
Analysis of copy number deletions. **(a)** Deletions affecting the *BMPR2* locus in 23 PAH cases. Genes are indicated in orange and labelled with their respective gene symbol. Deletions are drawn as blue boxes above the genome axis (grey) showing the genomic position on chromosome 2. The grey box highlights the location of *BMPR2*. **(b)** Locus zoom on *BMPR2* highlighting the focal deletions affecting one or more exons. **(c)** WGS coverage profiles of a selected set of smaller and larger deletions, visualised with the Integrative Genomics Viewer (IGV)^18^, with deletions highlighted by red bars. **(d)** and **(e)** Manhattan plots of the genome-wide case-control comparison of large deletions. In **(d)** all subject are considered. In **(e)** subject with larger deletions affecting the *BMPR2* locus are excluded. The adjusted *P* value threshold of 5 x 10^−8^ for genome-wide significance is indicated by the red line.

Of the other genes previously reported in PAH we identified deleterious heterozygous rare variants in *ACVRL1* (9 cases, 0.9%), *ENG* (6 cases, 0.6%), *SMAD9* (4 cases, 0.4%), *KCNK3* (4 cases, 0.4%), and *TBX4* (14 cases, 1.3%). We identified one case with highly deleterious variants in both *BMPR2* (p.Cys123Arg) and *SMAD9* (p.Arg294Ter). Details of consequence types, deleteriousness and conservation scores, and minor allele frequencies are provided in Supplementary Table 4. Fourteen cases (1.3%) with biallelic *EIF2AK4* mutations were found^19^. No pathogenic coding variants in *CAV1, SMAD1* or *SMAD4* were identified. Taken together, rare causal variation in non-*BMPR2* disease genes (*TBX4*, *ENG*, *ACVRL1*, *SMAD9*, *KCNK3* and *EIF2AK4*) accounted for 4.7% of the entire PAH cohort. The clinical characteristics of cases with variants in these previously reported genes are shown in Supplementary Table 5.

In a case-control comparison of the frequencies of deleterious variants confined to the previously reported PAH genes, we observed significant overrepresentation of rare variants in *BMPR2, TBX4*, *ACVRL1* and biallelic variants in *EIF2AK4* only (*P* < 0.05) (Supplementary Table 6).

### Identification of novel PAH disease genes

The strategy to identify novel causative genes in PAH employed a series of case-control analyses (Figure 1b). To identify signals that might be masked by variants in previously reported PAH genes, we excluded subjects with rare variants and deletions in *BMPR2*, *EIF2AK4*, *ENG*, *ACVRL1*, *TBX4*, *SMAD9* and *KCNK3*. A genome-wide comparison of protein truncating variants (PTVs), representative of high impact variants, identified a higher frequency of PTVs in *ATP13A3* (6 cases) (*P*_*adj*_ = 0.0346). Moreover, we identified additional PTVs in several putative PAH genes, including *EVI5* (5 cases, 1 control) and *KDR* (4 cases, 0 controls) (Figure 3a), that require further validation to evaluate their contribution to PAH pathogenesis (Supplementary Table 7).

**Figure 3.**
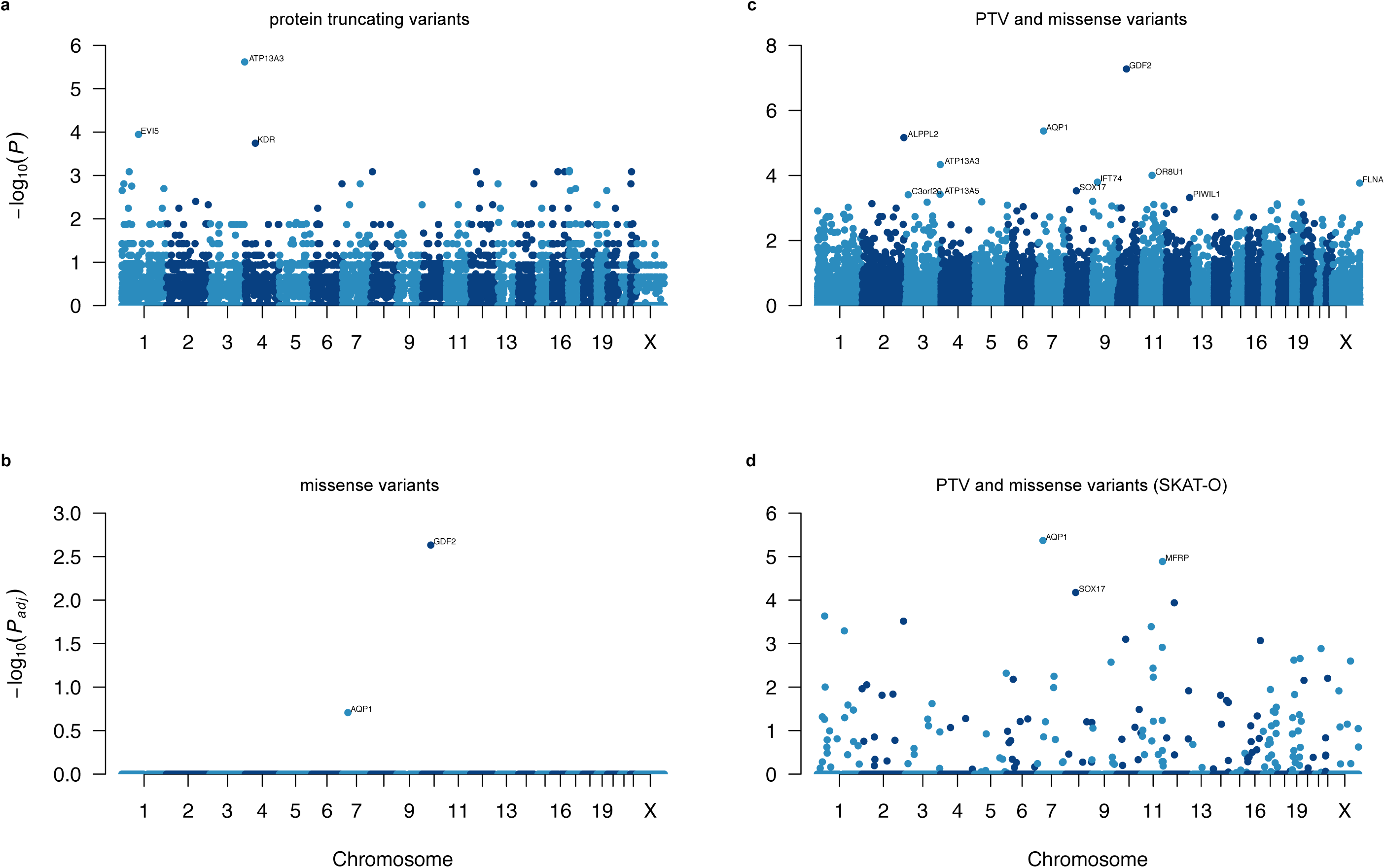
Manhattan plots of the rare variant analyses, having excluded cases carrying rare variants in previously established PAH genes. Filtered variants were grouped per gene. We tested for an excess of variants in PAH cases within genes using Fisher’s exact test. The negative decadic logarithm of unadjusted or adjusted *P*-values are plotted against the chromosomal location of each gene. **(a)** Burden test of rare PTVs. **(b)** Burden test of rare deleterious missense variants. **(c)** Burden test combining rare PTVs and likely deleterious missense variants. **(d)** SKAT-O test of rare PTVs and missense variants.

We next analysed rare missense variants overrepresented in the PAH cohort, again excluding subjects with variants in the previously reported PAH genes. This revealed significant overrepresentation of rare variants in *GDF2* after correction for multiple testing (*P_adj_* = 0.0023), followed by *AQP1* (Figure 3b and Supplementary Table 8). Next, in a combined analysis of rare missense variants and PTV, only *GDF2* remained significant (*P* = 0.001). Rare variants in additional putative genes occurred at higher frequency in cases compared to controls, including *AQP1, ALPPL2*, *ATP13A3*, *OR8U1*, *IFT74*, *FLNA*, *SOX17*, *ATP13A5*, *C3orf20* and *PIWIL1* (uncorrected *P* value < 0.0005), but were not significant after correction for multiple testing (Figure 3c and Supplementary Table 9).

In order to increase power to detect rare associations, we deployed SKAT-O on filtered rare PTVs and missense variants. Excluding previously reported genes, this analysis revealed an association with rare variants in *AQP1* (*P_adj_* = 4.28x10^−6^) and *SOX17* (*P_adj_* = 6.7x10^−5^) (Figure 3d). *AQP1* and *SOX17* were both also nominally significant in the combined burden tests, described above. Association was also found with rare variants in *MFRP* (*P_adj_* = 1.3x10^−5^). However, we consider *MFRP* a false-positive finding for reasons given in the Discussion. Supplementary Table 10 shows the top 50 most significant genes identified by SKAT-O, providing further candidates to be evaluated in future studies. Details of rare variants in novel PAH genes (*GDF2, ATP13A3, AQP1, SOX17*) identified in cases are provided in Supplementary Table 11.

Notably, a genome-wide assessment of larger structural variation did not identify any additional large deletions after exclusion of subjects harbouring deletions in *BMPR2* (Figure 2d-e).

The proportion of PAH cases with mutations in the new genes was 3.5%. The clinical characteristics of PAH cases with mutations in these genes are provided in Supplementary Table 5b. Of note, cases with mutations in *SOX17* and *AQP1* were significantly younger at diagnosis (32.8 ± 16.2 years [*P* = 0.002] and 36.9 ± 14.3 years [*P* = 0.013], respectively) compared to cases with no mutations in the previously established genes (51.7 ± 16.6 years).

### Independent validation and familial segregation analysis

To provide further validation of the potentially causal role of mutations in the new genes identified, we examined whole-exome data from an independent UK family with three affected individuals across two generations. Microsatellite genotyping across chromosome 2q33 had previously demonstrated non-sharing of haplotypes in affected individuals, consistent with exclusion of linkage to the *BMPR2* locus. No pathogenic variants were identified in the protein-coding regions of the *BMPR2* gene or other TGF-β pathway genes. Analysis of exome sequence data from individual II-1 identified a novel heterozygous c.411C>G (p.Y137*) PTV in the *SOX17* gene. Segregation analysis in the extended family demonstrated that the mutation had arisen *de novo* in the affected father (II-1) and was transmitted to the affected offspring (III-1). All unaffected family members were confirmed as wild-type (Figure 4a).

**Figure 4.**
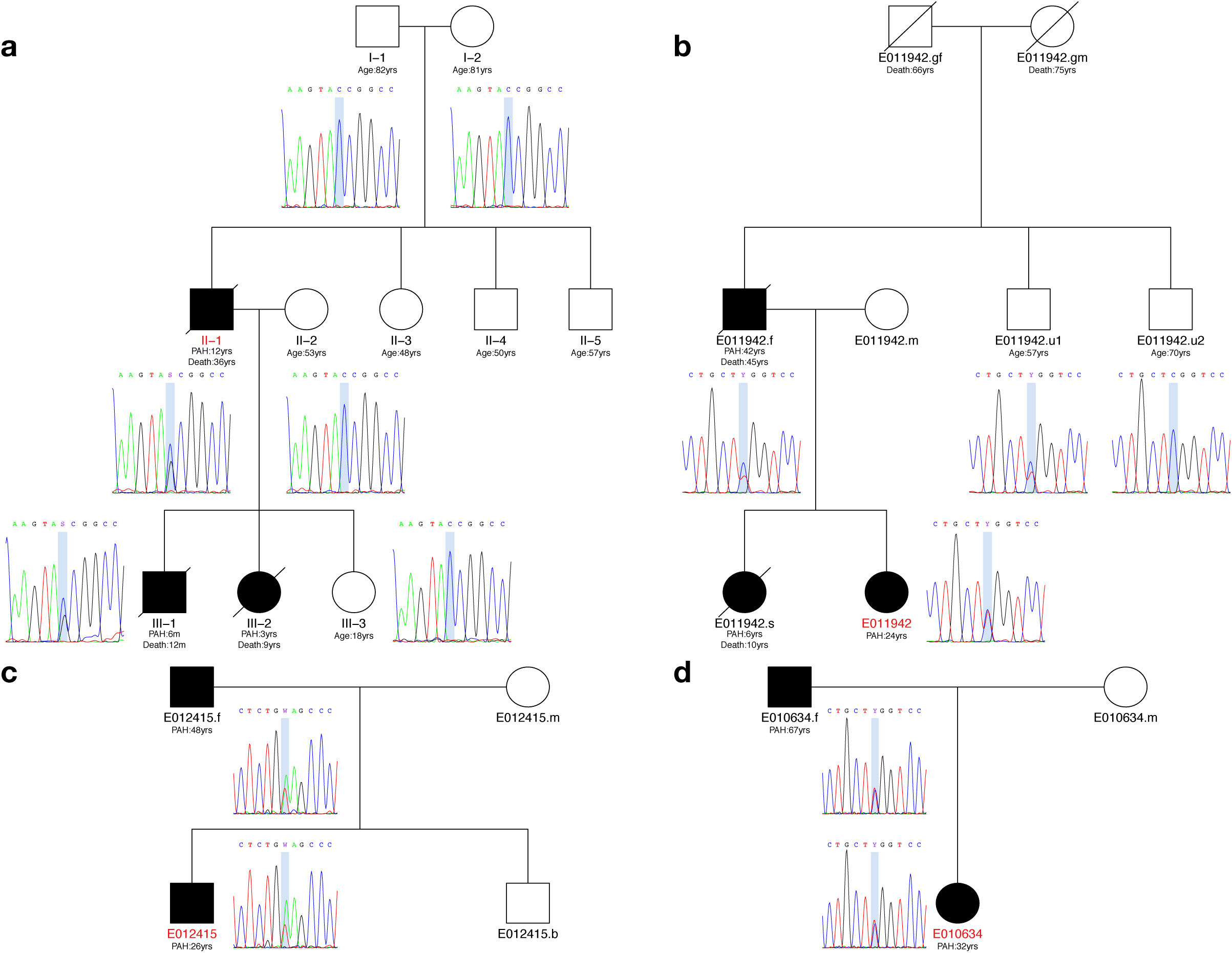
Pedigree structures and analysis of familial transmission of variants in *AQP1* and *SOX17*. **(a)** Individual II.1 harbours a heterozygous *de novo SOX17* c.411C>G (p.Y137*) PTV resulting in a premature termination codon, which has been transmitted to the affected male (III.1). No unaffected family members carry the variant. No sample was available from subject III.2. **(b)** Proband E011942 has inherited a heterozygous *AQP1* c.583C>T (p.R195W) missense variant from her affected father. No sample was available from the affected sister of the proband. The younger healthy uncle of the index case also carries the *AQP1* variant. No samples or further clinical information was available for the grandparents, who were not known to have cardiopulmonary disease. **(c)** Both the proband E012415 and her father are affected and carry the rare *AQP1* c.527T>A (p.V176E) missense variant. There was no further information available about the siblings of the father. **(d)** Subject E010634 has inherited the heterozygous *AQP1* c.583C>T (p.R195W) missense variant from her affected father. No rare variants in previously reported PAH genes were identified in any of theses families. Index cases are highlighted in red.

Three HPAH subjects harbouring rare variants in *AQP1*, identified in the NIHR BR-RD WGS study, were also selected for familial co-segregation analysis (Figure 4b-d). No pathogenic variants in any of the previously reported genes were identified in these families. The first pedigree comprised three affected individuals across two generations. Sanger sequencing confirmed the presence of the heterozygous *AQP1* c.583C>T (p.R195W) missense variant in the proband (E011942), the affected father (E011942.f) and the healthy younger paternal uncle (E011942.u1). An additional unaffected uncle did not carry the AQP1 variant. These results indicate likely incomplete penetrance in the unaffected carrier, as observed in *BMPR2* families^20^. No additional clinical information was available for the deceased grandparents (Figure 4b). The remaining two families comprised affected parent-offspring individuals. By Sanger sequencing we independently confirmed a heterozygous *AQP1* c.527T>A (p.Val176Glu) missense variant in proband (E012415) and his affected father (Figure 4c), as well as a heterozygous *AQP1* c.583C>T (p.R195W) missense variant in proband (E010634) and her affected father (Figure 4d). These results highlight recurrent *AQP1* variation across unrelated families and demonstrate co-segregation with the phenotype.

### Predicted functional impact of variants in novel PAH genes

To evaluate the potential functional impact of rare variants identified in the likely causative new genes we performed structural analysis of *GDF2*, *ATP13A3*, *AQP1*, and *SOX17*. In addition we undertook a functional analysis of the *GDF2* variants identified.

Heterozygous mutations in *GDF2* exclusive to PAH cases comprised 1 frameshift variant and 7 missense variants. *GDF2* encodes growth and differentiation factor 2, also known as bone morphogenetic protein 9 (BMP9), the major circulating ligand for the endothelial BMPR2/ACVRL1 receptor complex^21^. Amino acid substitutions were assessed against the published crystal structure^22^ of the prodomain bound form of GDF2 (Figure 5). Variants clustered at the interface between the prodomain and growth factor domain. Since the prodomain is important for the processing of GDF2, it is likely that amino acid substitutions reduce the stability of the prodomain-growth factor interface. In keeping with these predictions, HEK293T cells transfected with *GDF2* variants exclusive to PAH cases, demonstrated reduced secretion of mature GDF2 into the cell supernatants (Figure 5d), compared with wild type GDF2.

**Figure 5.**
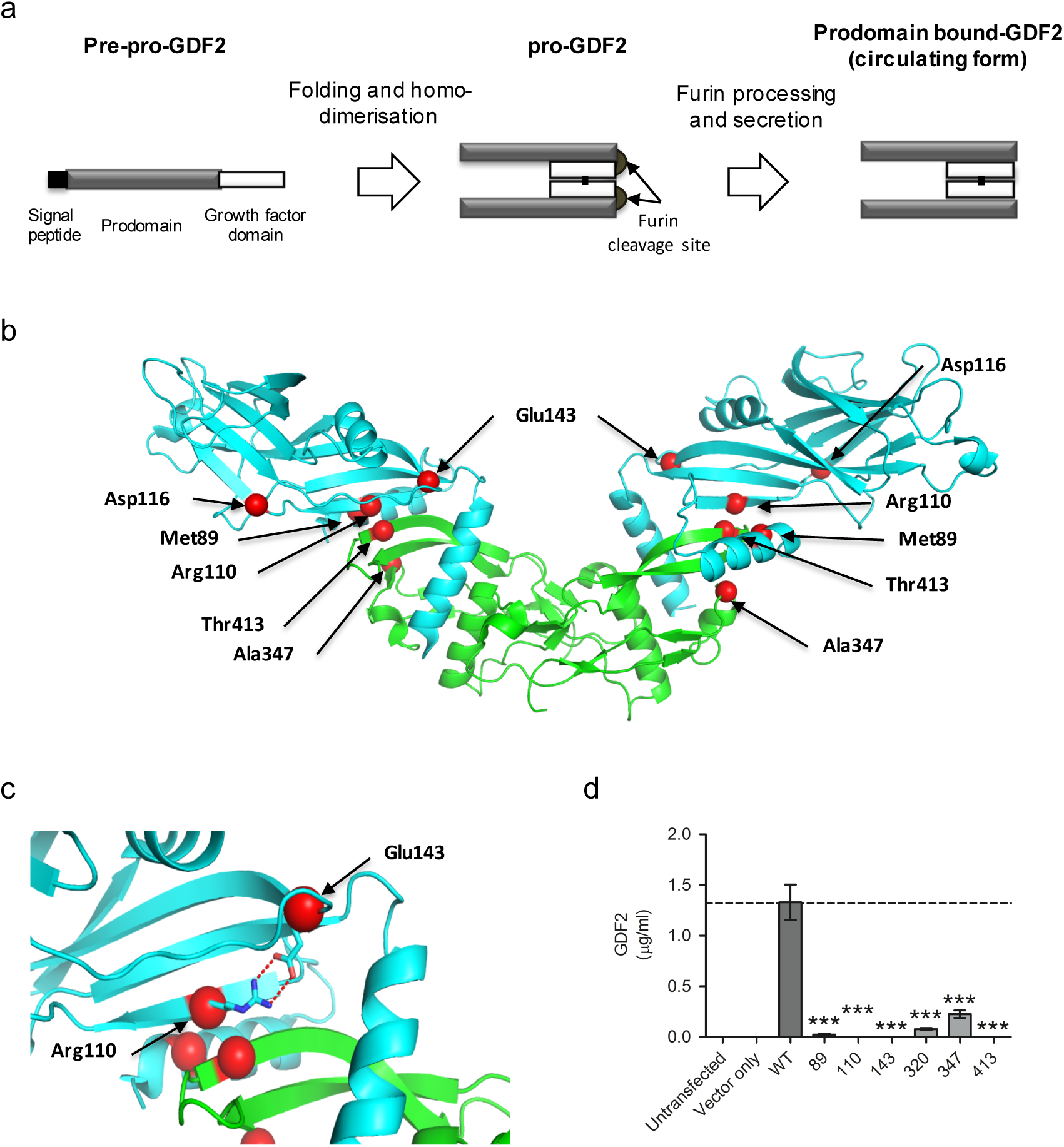
Structural analysis of *GDF2* mutations. **(a)** Schematic diagram of GDF2 processing. The pre-pro-protein is processed into the mature growth factor domain (GFD) bound to the prodomain upon secretion^23^. **(b)** Plot of *GDF2* mutations found only in PAH cases superimposed on the structure of prodomain bound GDF2. (pdb code 4YCG)^22^. The GDF2 growth factor domain is shown in green and the prodomain in cyan. **(c)** Magnified view of the Arg110 and Glu143 mutations. The wild type amino acids make double salt bridges to stabilise the prodomain conformation at the interface between the growth factor domain and prodomain. The E143K and R110W mutations both disrupt these interactions, destabilising the interaction between the growth factor domain and prodomain. **(d)** GDF2 levels secreted into supernatants of HEK293T cells transfected with likely pathogenic variants found in PAH cases, compared with wild type GDF2 and cells transfected with an empty vector. *** *P* < 0.001 by ANOVA.

We identified 3 heterozygous frameshift variants, 2 stop gained, 2 splice region variants in *ATP13A3*, which are predicted to lead to loss of ATPase catalytic activity (Figure 6a). In addition, we identified 4 heterozygous likely pathogenic missense variants in PAH cases, two near the conserved ATPase catalytic site and predicted to destabilise the conformation of the catalytic domain (Figure 6b-d). The distribution of variants (Figure 6a) suggests that these mutations impact critically on the function of the protein.

**Figure 6.**
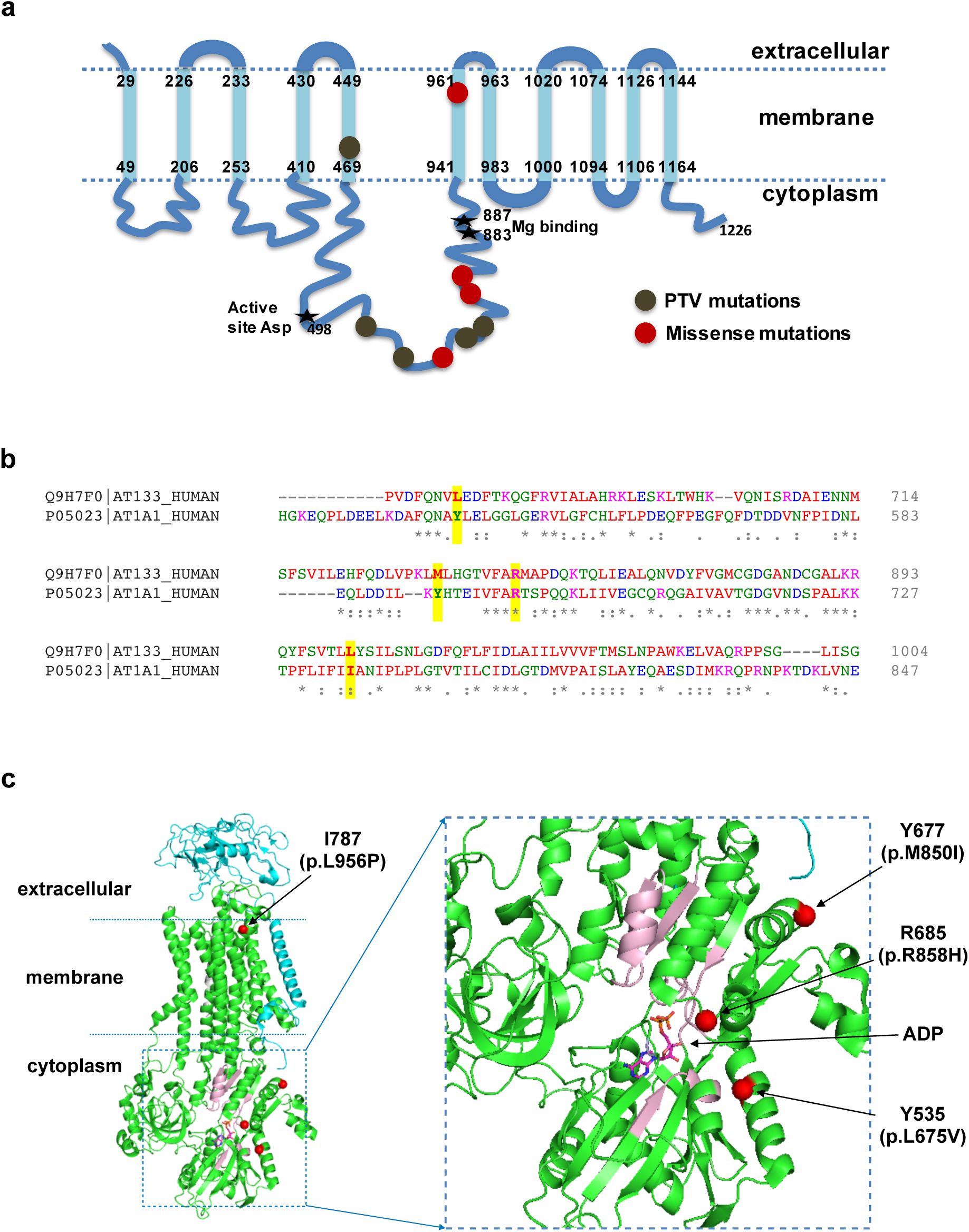
Structural analysis of *ATP13A3* mutations. **(a)** Topology of ATP13A3, plotted according to UniProtKB Q9H7F0. Frameshift and stop-gained mutations identified in PAH cases are shown as khaki circles, and missense mutations as red circles. Frameshift/stop-gained mutations are predicted to truncate the protein prior to the catalytic domain and essential Mg binding sites, leading to loss of ATPase activity. **(b)** Sequence alignment of ATP13A3 with ATP1A1 (P05024), of which the high resolution structure was used for the structural analysis in **(c)**. The conserved regions of ATP13A3 and ATP1A1, essential for ATPase activity^24^, show good alignment (data not shown). Only regions containing the missense PAH mutations are shown, with positions of the four missense mutations highlighted in yellow above the sequences. **(c)** Structural analysis of the 4 PAH missense mutations plotted on the ATP1A1 crystal structure based on the sequence alignment in **(b)** (pdb code 3wgu)^25^. Green: **a** subunit (P05024), cyan: **(3** subunit (P05027), grey: y-subunit transcript variant a (Q58k79). Y535, Y677, R685 and I787 are the numbering in ATP1A1. Positions of the four missense mutations found in PAH are labeled and highlighted by red circles. **(d)** Magnified view of the cytoplasmic region of the ATPase, showing the presence of ADP at the active site. The conserved regions essential for ATPase activity are shown in light pink. The L675V and R858H mutations are located close to the ATP catalytic region.

The majority of rare variants identified in *AQP1*, which encodes aquaporin 1, are situated within the critical water channel (Figure 7). In particular the p.Arg195Trp variant, identified in 5 PAH cases, locates at the hydrophilic face of the pore. This arginine at position 195 helps define the constriction region of the AQP1 pore structure and is conserved across the water specific aquaporins^26^. Rare variants in *SOX17*, included 4 nonsense variants (including the PTV identified in the additional UK family) predicted to lead to loss of the beta-catenin binding region, and 6 missense variants predicted to disrupt interactions with Oct4 and beta-catenin^27, 28^ (Figure 8).

**Figure 7.**
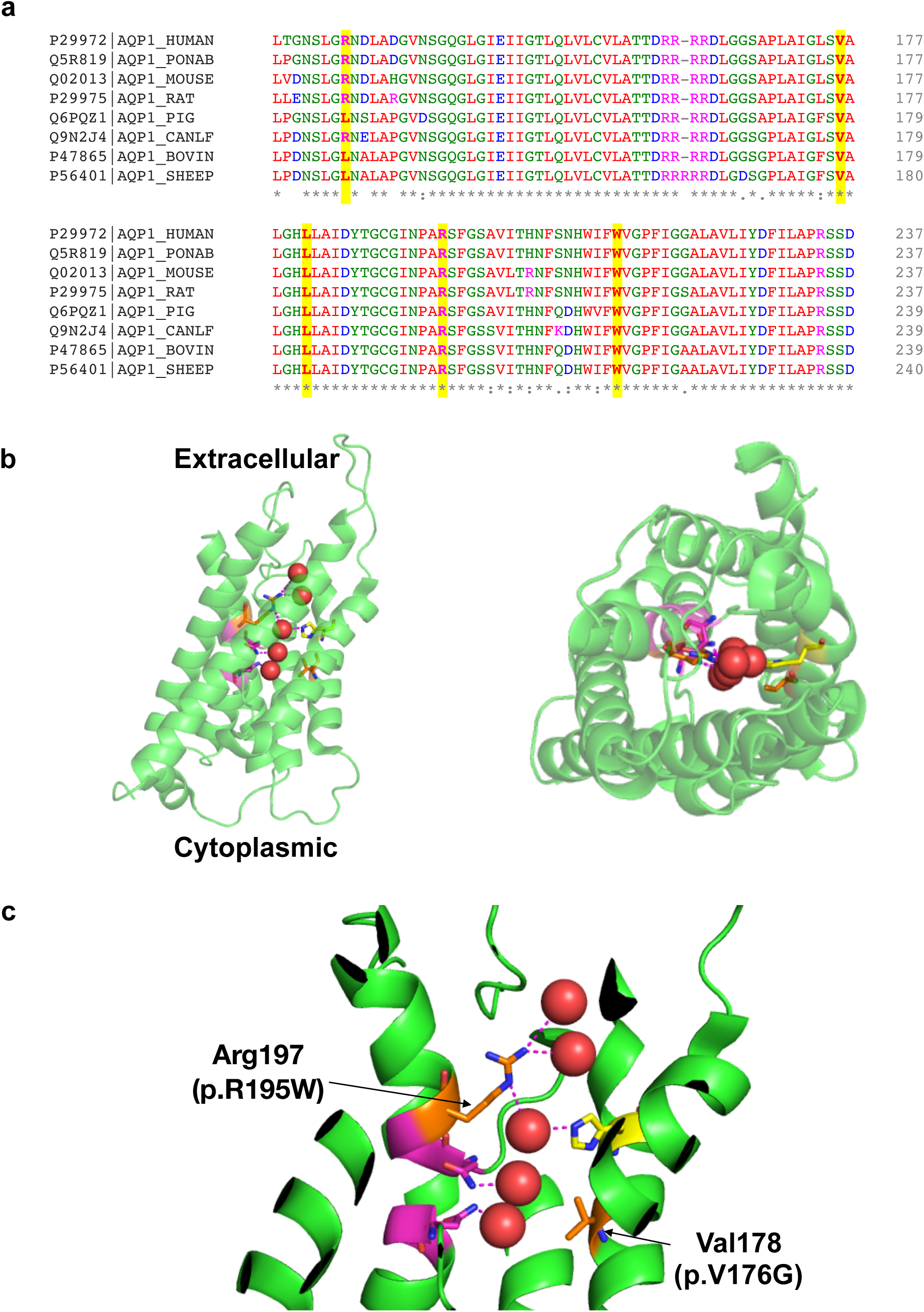
Structural analysis of *AQP1* mutations. **(a)** Multiple sequence alignment of human AQP1 with seven other mammals. The bovine AQP1 has the high resolution (2.2Å) published structure. Mutations identified in PAH cases are highly conserved and highlighted in yellow. **(b)** Crystal structure of bovine AQP1 (pdb code 1j4n)^26^. Left: side view; right: top view from the extracellular direction. AQP1 is shown as a semi-transparent cartoon and five water molecules in the water channel are shown as red spheres. Key residues lining the water channels are represented with stick structures. **(c)** Magnified view of the water channel, with H-bonds connected to water molecules in the channel highlighted. Two asparagine-proline-alanine (NPA) motifs, essential for the water transporting function of AQP1, are shown in magenta. Conserved His180 that constricts the water channel is shown in yellow. Mutations found in PAH cases, Arg195Trp and Val176Glu, are labelled and shown as orange stick structures. Arg195 and His180 are highly conserved in the known water channels and are strong indicators of water channel specificity. Arg195Trp and Val176Glu mutations are predicted to disrupt the conformation of this conserved water channel.

**Figure 8.**
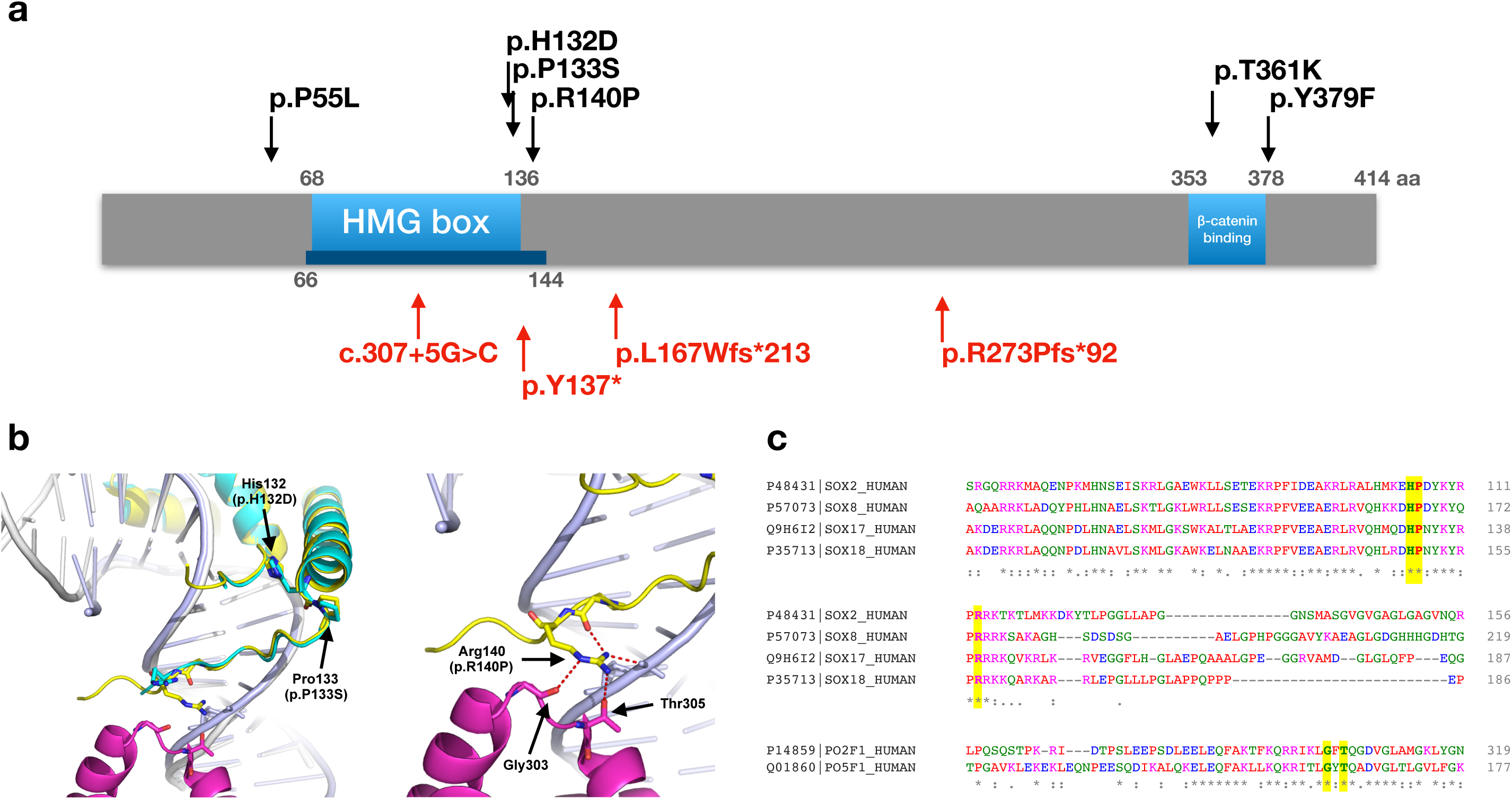
Structural analysis of *SOX17* mutations. **(a)** Schematic diagram of human SOX17 (Q9H6I2), based the UniProtKB annotation, and published reports^27^. Red arrows indicate PTVs and black arrows indicate missense mutations identified in PAH patients. The blue bar illustrates the region that is covered in the crystal structure (pdb code 3F27)^29^. The ability of SOX17 to activate transcription of target genes correlates with binding to β-catenin^27^. As illustrated, all PTVs lead to a loss of the β-catenin binding region. Two missense mutations are located within and very close to the minimum β-catenin binding regions, and both are highly conserved, indicating they are likely to be important for β-catenin binding. **(b)** Structural analysis of HMG domain missense mutations found in PAH patients. Left, Superposition of SOX17/DNA structure (pdb code 3F27, Sox17: cyan, DNA: grey)^29^ onto SOX2/DNA/Oct1 structure (pdb code 1GT0, Sox2: yellow, Oct1: magenta, DNA: light blue)^28^. Right: Magnified view of the interactions around Arg140 in the SOX2/DNA/Oct structure. Arg140 in SOX2 makes multiple H-bond interactions and mutating this Arg in SOX2 abolishes the interaction with transcription factors Pax6 and Oct4^28^. SOX2 and SOX17 both bind to Oct4^30^ and SOX17 K122E mutant can replace SOX2 in maintaining stem cell pluripotency^30^, indicating this region in SOX17 may interact with Oct4, similar to SOX2. The three missense mutations in *SOX17* will likely disrupt interaction with Oct4. **(c)** Supporting the analysis in **(b)**, sequence alignment shows that the HMG domain of SOX2 and SOX17 as well as SOX8 (P57073) and SOX18 (P35713) share high sequence identity and the three mutations found in PAH (highlighted in yellow) are highly conserved emphasising their functional importance. Similarly, the Gly and Thr that interact with Arg140 in SOX2 (highlighted in yellow) are also conserved between Oct1 (PO2F1) and Oct4 (PO5F1).

GDF2 is known to be secreted from the liver, but the cellular localization of proteins encoded by the other novel genes is less well characterised. Thus we employed immunohistochemistry to examine localization in the normal and hypertensive human pulmonary vasculature. Figure 9 shows that AQP1, ATP13A3 and SOX17 are predominantly localised to the pulmonary endothelium in normal human lung and to endothelial cells within plexiform lesions of patients with idiopathic PAH. In addition, we determined the relative mRNA expression levels of *AQP1*, *ATP13A3* and *SOX17* in primary cultures of pulmonary artery smooth muscle cells (PASMCs), pulmonary artery endothelial cells (PAECs) and blood outgrowth endothelial cells (BOECs)^31^. AQP1 was expressed in PASMCs and endothelial cells, with a trend towards higher levels in PASMCs (Figure 10a). ATP13A3 was highly expressed in both cell types (Figure 10b), whereas SOX17 was almost exclusively expressed in endothelial cells (Figure 10c). Although AQP1 and SOX17 are known to play roles in endothelial function, the function of ATP13A3 in vascular cells is entirely unknown. Thus, we determined the impact of ATP13A3 knockdown on proliferation and apoptosis of BOECs. Loss of ATP13A3 led to marked inhibition of serum-stimulated proliferation of BOECs, and increased apoptosis in serum-deprived conditions (Figure 10d-f).

**Figure 9.**
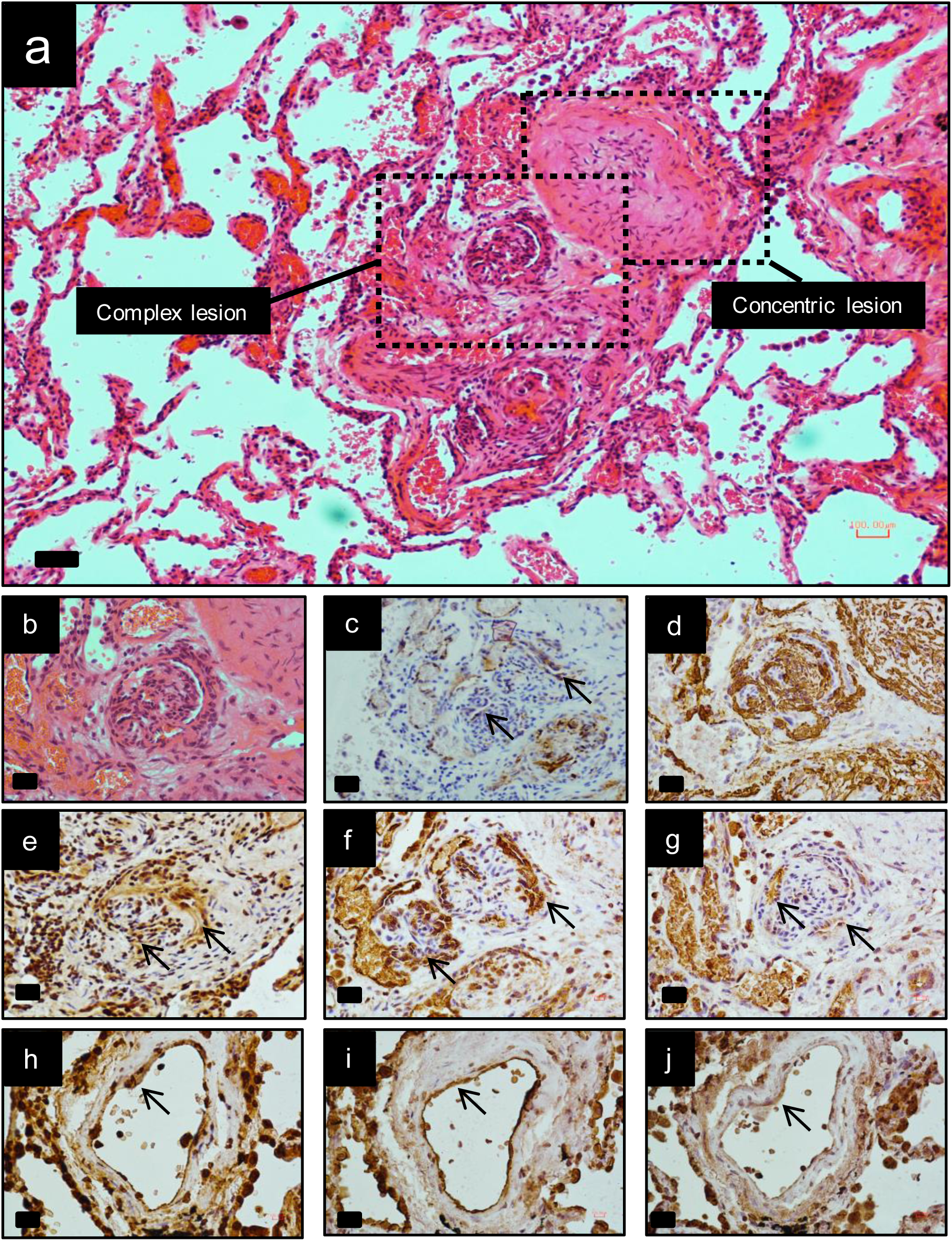
Immunolocalisation of AQP1, ATP13A3 and SOX17 in normal and PAH lung. The typical histological findings (haematoxylin and eosin staining) of concentric vascular lesions with associated plexiform lesions are shown (**a**). Higher magnification images of plexiform lesion (**b**), with frequent endothelialised channels (**c**; anti-CD31) surrounded by myofibroblasts (**d**; anti-SMα). Additional high magnification images demonstrating endothelial expression of ATP13A3 (**e**), AQP1 (**f**) and SOX17 (**g**) in PAH lung. Controls lung sections demonstrating predominantly endothelial expression of ATP13A3 (**h**), AQP1 (**i**) and SOX17 (**j**). (Scale bars = 50μm).

**Figure 10.**
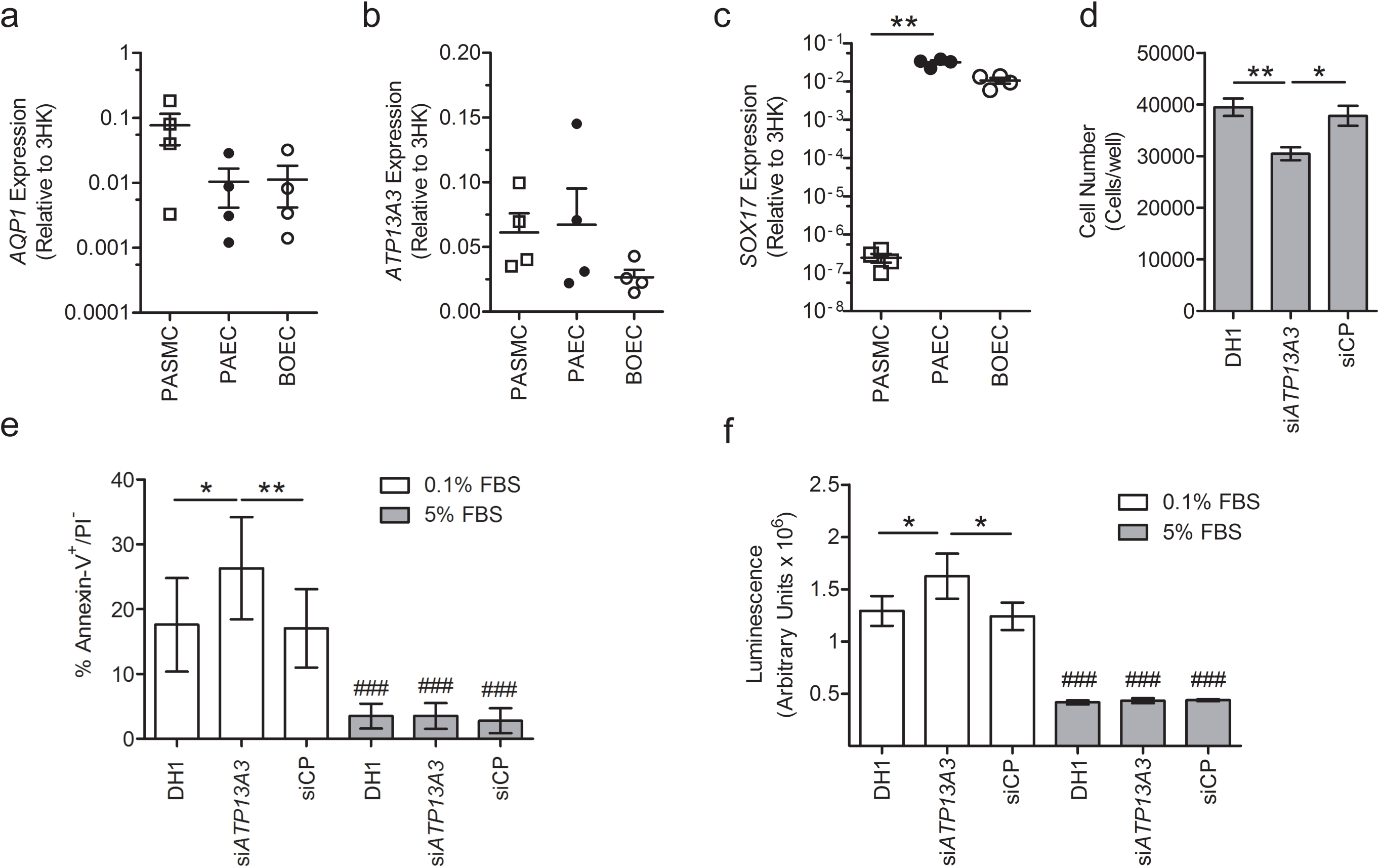
Functional studies of novel genes. (**a**-**c**). Expression of (**a**) *AQP1*, (**b**) *ATP13A3* and (**c**) *SOX17* mRNA in human pulmonary artery smooth muscle cells, pulmonary artery endothelial cells and blood outgrowth endothelial cells (BOECs) (n=4 biological replicates of each). Relative expression of each transcript was normalised to three reference genes, *ACTB*, *B2M* and *HPRT*. (**d**) Proliferation of BOECs in 5% FBS over 6 days. Cells were transfected with DharmaFECT1 alone (DH1), si*ATP13A3* or non-targeting siRNA control (siCP) (**e-f**) Quantification of apoptosis in BOECs, defined as Annexin V+/PI-cells, in BOECs transfected with si*ATP13A3* or siCP in complex with DH1 followed by 24hr treatment with 0.1% FBS or 5% FBS (n=4 biological repeats). (**f**) Measurement of apoptosis via Caspase-Glo 3/7 activity measurements in BOECs transfected with si*ATP13A3* or siCP in complex with DH1, followed by 16hr treatment in 0.1% FBS or 5% FBS. Data are from a single experiment (n=4 wells) representative of 3 biological repeats. Data were analysed using a One-way analysis of variance with post hoc Tukey’s test for multiple comparisons in **d** and **f**. Data were analysed using a repeated measures One-way analysis of variance with post hoc Tukey’s for multiple comparisons in **e**. *P<0.05, **P<0.01 within treatment groups. ^###^P<0.001 for effect of ligand against control for same transfection condition.

## Discussion

We report the most comprehensive analysis to date of rare genetic variation in a large cohort of index cases with idiopathic and heritable forms of PAH. Whilst we utilised WGS, the main goal was the identification of novel rare causal variation underlying PAH in the protein coding sequence. The approach involved a rigorous case-control comparison using a tiered search for variants. First, we searched for high impact PTVs overrepresented in cases, having excluded previously established PAH genes. This revealed PTVs in *ATP13A3*, a poorly characterised P-type ATPase of the P5 subfamily^32^. There is little information regarding the function of the ATPase, *ATP13A*3, which appears widely expressed in mouse tissues^32^. Although, the precise substrate specificity is unknown, ATP13A3 plays a role in polyamine transport^33^. Based on available RNA sequencing data, *ATP13A3* is highly expressed in human pulmonary vascular cells and cardiac tissue (https://www.encodeproject.org). We confirmed that *ATP13A3* mRNA is expressed in primary cultured pulmonary artery smooth muscle cells and endothelial cells, and provide preliminary data that loss of *ATP13A3* inhibits proliferation and increases apoptosis of endothelial cells. These findings are consistent with the widely accepted paradigm that endothelial apoptosis is a major trigger for the initiation of PAH^34, 35^. It will be of considerable interest to determine the role of *ATP13A3* in vascular cells and whether it is functionally associated with BMP signalling, or represents a distinct therapeutic target in PAH.

Analysis of missense variation, and a combined analysis of all predicted deleterious variation, revealed that mutation at the *GDF2* gene is also significant determinant of predisposition to PAH. Of the new genes identified, *GDF2* provides further evidence for the central role of the BMP signalling pathway in PAH. *GDF2* encodes the major circulating ligand for the endothelial BMPR2/ACVRL1 receptor complex^21^. Taken together, the genetic findings suggest that a deficiency in GDF2/BMPR2/ACVRL1 signalling in pulmonary artery endothelial cells is critical in PAH pathobiology. The majority of *GDF2* variants detected in our adult-onset PAH cohort were heterozygous missense variants, in contrast to a previous case report of childhood onset PAH due to a homozygous nonsense mutation^36^. The finding of causal *GDF2* variants in PAH cases, associated with reduced production of GDF2 from cells, provides further support for investigating replacement of this factor as a therapeutic strategy in PAH^37^.

To maximise the assessment of rare variation in a case-control study design, we deployed the SKAT-O test. This approach revealed a significant association of rare variation in the aquaporin gene, *AQP1*, and the transcription factor encoded by *SOX17*. Of note, both *AQP1* and *SOX17* were within the top 8 ranked genes in our combined PTV and missense burden test analysis (Supplementary Table 9), providing further confidence in their causative contribution to PAH.

Aquaporin 1 belongs to a family of membrane proteins that function as channels facilitating water transport in response to osmotic gradients^26^, and AQP1 is known to promote endothelial cell migration and angiogenesis^38^. Thus, approaches that maintain or restore pulmonary endothelial function could offer new therapeutic directions in PAH. Conversely, AQP1 inhibition in pulmonary artery smooth muscle cells ameliorated hypoxia-induced pulmonary hypertension in mice^39^, suggesting that further studies are required to determine the key cell type impacted by *AQP1* mutations in human PAH, and the functional impact of these *AQP1* variants on water transport. The demonstration of familial segregation of *AQP1* variants with PAH provides further support for the potentially causal role of these mutations in disease. However, we also identified an unaffected *AQP1* variant carrier consistent with reduced penetrance, which is well described for other PAH genes, including *BMPR2*.

Although functional studies are required to confirm the mechanisms by which mutations in *SOX17* cause PAH, this finding provides additional support for the vascular endothelium as the major initiating cell type in this disorder. *SOX17* encodes the SRY-box containing transcription factor 17, which plays a fundamental role in angiogenesis^40^ and arteriovenous differentiation^41^. Moreover, conditional deletion of *SOX17* in mesenchymal progenitors leads to impaired formation of lung microvessels^42^. The demonstration of familial segregation of the *SOX17* p.Y137* PTV with early onset PAH provides additional evidence for a causal role for these variants in PAH. The co-existence of a patent ductus arteriosus in the index case and an atrial septal defect (ASD) in one of the affected offspring is of interest and suggests an association with congenital heart disease. Small ASDs are not uncommon in idiopathic PAH, and a more detailed clinical phenotyping of SOX17 mutation carriers will be required to determine whether the presence of ASDs and other congenital heart abnormalities are more common in carriers of these mutations.

Whilst the SKAT-O analysis also provided support for the *MFRP* gene, recessive bi-allelic mutations in *MFRP* cause retinal degeneration and posterior microphthalmos^43^. The expression of *MFRP* transcripts is largely confined to the central nervous system^44^ and the majority of variants were present in the Genome Aggregation Database (GnomAD, http://gnomad.broadinstitute.org). On the basis of these considerations, variants in *MFRP* are unlikely to contribute to PAH aetiology.

This analysis provides new insights on the frequency and validity of previously reported genes in PAH. We confirmed that mutations in *BMPR2* are the most common genetic cause and validated rare causal variants in *ACVRL1*, *ENG*, *SMAD9*, *TBX4*, *KCNK3* and *EIF2AK4*. Although our findings question the validity of *CAV1*, *SMAD1* and *SMAD4* as causal genes, previous reports might represent private mutations occurring in very rare families. The use of WGS in this study allowed closer interrogation of larger deletions around the *BMPR2* locus than has been possible previously. Nevertheless, additional analyses are required to determine the full impact of structural variation (inversions, duplications, smaller deletions) at this and other loci.

The non-PAH cohort used in the case-control comparisons for this study comprised individuals, or relatives of individuals, with other rare diseases recruited to the NIHR Bioresource for Rare Diseases (NIHR BR-RD) in the UK (see Methods). In general, for very rare causal variants, the comparison between PAH cases and non-PAH rare disease controls should not reduce our ability to detect overrepresentation of rare variants in a particular gene in the PAH cohort, if mutations in that gene are specific to PAH. However, if rare variants in a gene were responsible for more than one phenotype, it is possible that this would reduce the power to detect overrepresentation in the PAH cohort. For example, if mutations occurred in different functional domains of the expressed protein, this might lead to PAH if mutations affected one domain, but other phenotypes if they affected another domain. Overcoming this potential limitation will require additional analysis of the functional impact of variants and their distribution within a gene, and more detailed information on the phenotypes of subjects in the non-PAH group.

Taken together, this study identifies rare sequence variation in new genes underlying heritable forms of PAH, and provides a unique resource for future large-scale discovery efforts in this disorder. Mutations in previously established genes accounted for 19.9% of PAH cases. Including new genes identified in this study (*GDF2, ATP13A3, AQP1, SOX17*), the total proportion of cases explained by mutations increased to 23.5%. It is likely that independent confirmation of the expanded list of putative genes identified in this study will increase further the proportion of cases explained by mutations, but this will require larger international collaborations. The results suggest that the genetic architecture of PAH, beyond mutations in *BMPR2*, is characterised by substantial genetic heterogeneity and consists of rare heterozygous coding region mutations shared by small numbers of cases. The contribution of rare variation within non-coding regulatory regions to PAH aetiology remains to be determined. This will require functional annotation of regulatory and other non-coding regions specific for relevant cell types, further case-control analyses of these regions and ultimately functional studies of gene regulation to assess the pathogenicity of non-coding variants. Our findings to date provide support for a central role of the pulmonary vascular endothelium in disease pathogenesis, and suggest new mechanisms that could be exploited therapeutically in this life-limiting disease.

## Methods

### Ethics and patient selection

Cases were recruited from the UK National Pulmonary Hypertension Centres, Universite Sud Paris (France), the VU University Medical Center Amsterdam (The Netherlands), the Universities of Gießen and Marburg (Germany), San Matteo Hospital, Pavia (Italy), and Medical University of Graz (Austria). All patients provided written informed consent (UK Research Ethics Committee: 13/EE/0325), or local forms consenting to genetic testing in deceased patients and non-UK cases. All cases had a clinical diagnosis of idiopathic PAH, heritable PAH, drug-and toxin-associated PAH, or PVOD/PCH established by their expert centre. An additional UK family diagnosed with HPAH was ascertained as described previously^45^. Blood and saliva samples were collected under written informed consent of the participants or their parents for use in gene identification studies. The non-PAH cohort for the case-control comparison comprised 6385 unrelated subjects recruited to the NIHR BR-RD study.

### Composition of non-PAH control cohort

The non-PAH control cohort consisted of subjects with bleeding, thrombotic and platelet disorders (15.5%), cerebral small vessel disease (2.1%), Ehlers-Danlos syndrome (0.3%), subjects recruited to Genomics England Ltd (19.8%), hypertrophic cardiomyopathy (3.6%), intrahepatic cholestasis of pregnancy (4.1%), Leber hereditary optic neuropathy (0.9%), multiple primary tumours (7.8%), neuropathic pain disorder (2.6%), primary immune disorders (15.3%), primary membranoproliferative glomerulonephritis (2.3%), retinal dystrophies/paediatric neurology and metabolic disease (19.8%), stem cell and myeloid disorders (2.1%), steroid resistant nephrotic syndrome (3.6%), and others (0.3%), or their first degree relatives.

### High-throughput sequencing

DNA extracted from venous blood underwent whole genome sequencing using the Illumina TruSeq DNA PCR-Free Sample Preparation kit (Illumina Inc., San Diego, CA, USA) and Illumina HiSeq 2000 or HiSeq X sequencer, generating 100 - 150 bp reads with a minimum coverage of 15X for ~95% of the genome (mean coverage of 35X). Whole-exome sequencing was conducted for individual II-1 (Figure 4a) using genomic DNA extracted from peripheral blood. Paired-end sequence reads were generated on an Illumina HiSeq 2000.

### Data pre-processing and formation of analysis-ready data sets

Sequencing reads were pre-processed by Illumina with Isaac Aligner and Variant Caller (v2, Illumina Inc.) using human genome assembly GRCh37 as reference. Variants were normalised, merged into multi-sample VCF files by chromosome using the gVCF aggregation tool agg (https://github.com/Illumina/agg) and annotated with Ensembl’s Variant Effect Predictor (VEP). Following read alignment to the reference genome (GRCh37), variant calling and annotation of whole-exome data for individual II:1 were performed using GATK UnifiedGenotyper^46^ and ANNOVAR^47^, respectively. Annotations included minor allele frequencies from other control data sets (i.e. ExAC^48^, 1000 Genomes Project^49^ and UK10K^50^) as well as deleteriousness and conservation scores (i.e. CADD^51^, SIFT^52^, PolyPhen-2^53^ and Gerp^54^) enabling further filtering and assessment of the likely pathogenicity of variants(. To take forward only high quality calls, the pass frequency (proportion of samples containing alternate alleles that passed the original variant filtering) and call rate (proportion of samples with reference or alternate genotypes) were combined into the overall pass rate (OPR: pass frequency x call rate) and variants with an OPR of 80% or higher were retained.

### Estimation of ethnicity and relatedness

We estimated the population structure and relatedness based on a representative set of SNPs using the R package GENESIS to perform PC-Air^55^ and PC-Relate^56^, respectively. The selected 35,114 autosomal SNPs were present on Illumina genotyping arrays (HumanCoreExome-12v1.1, HumanCoreExome-24v1.0, HumanOmni2.5-8v1.1), do not overlap quality control excluded regions or multiallelic sites in the 1000 Genomes (1000G) Phase 3 dataset^49^, do not have any missing genotypes in NIHR BR-RD, had a MAF of 0.3 or above and LD pruning was performed using PLINK^57^ with a window size of 50 bp, window shift of 5 bp and a variance inflation factor threshold of 2. The 2,110 samples from the 1000G Project including the European (EUR), African (AFR), South Asian (SAS) and East Asian (EAS) populations (excluding the admixed American population) were filtered for the selected SNPs and the filtered data were used to perform a principal component analysis (PCA) using PC-Air. We modelled the scores of the leading five principal components as data generated by a population specific multivariate Gaussian distribution and estimated the corresponding mean and covariance parameters. Genotypes from the NIHR BR-RD samples were projected onto the loadings for the leading five principal components from the 1000G PCA and we computed the likelihood that each sample belonged to each subpopulation under a mixture of multivariate Gaussians models. Each sample was allocated to the population with the highest likelihood, unless the highest likelihood was similar to likelihood values for other populations, as might be expected for example under admixed ancestry or if the sample came from a population not included in 1000G. Such ambiguous samples were labeled as “other”. PC-Relate was used to to identify related individuals in NIHR BR-RD. We used the first 20 PCs from PC-Air to adjust for relatedness and extracted the pairwise Identity-By-State distances and kinship values. The pairwise information was used by Primus to infer family networks and calculate the maximum set of unrelated samples.

Of the 9,110 NIHR BR-RD samples, we assigned 80.2% to Non-Finish European (n=7,307), 7.2% to South Asian (n=649), 2.3% to African (n=213), 0.08% to East Asian (n=78), 0.02% to Finnish-European (n=19) and 9.2% to Other (n=844) and retrieved a maximum set of 7,493 unrelated individuals (UWGS10K), representing 82.2% of the entire NIHR BR-RD cohort.

### Cohort definition and allele frequency calculation

Based on the relatedness analysis, we defined the following sample subsets: (a) the maximum number of unrelated non-PAH controls (UPAHC, n=6385), (b) all affected PAH cases (PAHAFF, n=1048), and (c) all unrelated PAH index cases (PAHIDX, n=1038). These subsets were used to annotate the variants in the multi-sample VCF file with calculated minor allele frequencies using the fill-tags extension of BCFtools^58^.

### Rare variant filtering

Filtering of rare variants was performed as follows: 1) variants with a MAF less than 1 in 10,000 in UPAHC subjects, UK10K and ExAC were retained (adjusted for X chromosome variants to 1 in 8,000); 2) variants with a combined annotation dependent depletion deleteriousness (CADD) score of less than 15 were excluded. CADD scores were calculated using the CADD web service (http://cadd.gs.washington.edu) for variants lacking a score; 3) premature truncating variants (PTVs) or missense variants of the canonical transcript were retained; 4) missense variants predicted to be both tolerated and benign by SIFT and PolyPhen-2, respectively, were removed.

To identify likely causative mutations (as reported in Supplementary Table 5), variants in previously reported and putative genes, identified in this study, were examined in more detail to exclude variants that did not segregate in families (where data available). Furthermore, variants shared between cases and non-PAH controls, as well as variants of uncertain significance that co-occurred with previously reported causative mutations or high impact PTVs were also excluded.

### Burden analysis of protein-truncating and missense variants

Filtered variants were grouped per gene and consequence type (predicted PTV / missense) and subjects with at least one variant were counted (no double counting) per group and tested for association with disease. We applied a one-tailed Fisher’s exact test with *post hoc* Bonferroni correction to calculate the *P* value for genome-wide significance.

### Rare variant analysis using SKAT-O

To further investigate the aggregated effect that rare variants contribute to PAH aetiology, we applied a Sequence Kernel Association test (SKAT-O). SKAT-O increases the power of discovery under different inheritance models by combining variance-component and burden tests. Variants were filtered based on MAF as specified above, and only PTV and missense variants were included. For the analysis we implemented SKAT-O in RvTests v1.9.9^59^ with default parameters and weights being Beta(1,25), and applying a correction for read length, gender and the first five principal components of the ethnicity PCA. Variants were collapsed considering only the protein-coding region in the canonical transcript of the protein-coding genes in the genome assembly GRCh37.

### Analysis of large deletions

Copy number variation was identified using Canvas^60^ and Manta^61^. Deletions called by both Manta and Canvas with a reciprocal overlap of ≥ 20% were retained. Of these, deletions were excluded if both failed standard Illumina quality metrics or overlapped with known benign deletions in healthy cohorts^62^. Deletions with a reciprocal overlap of ≥ 50% between samples were merged and filtered for a frequency of less than 1 in 1,000 in WGS10K and overlapping exonic regions of protein coding genes (GRCh37 genome assembly). The number of subjects with deletions were added up by gene (no double counting of subjects) and tested for association with the disease. We applied a one-tailed (greater) Fisher’s exact test with Bonferroni *post hoc* correction for multiple testing to determine the *P* values for genome-wide significance.

### Confirmation of variants

Variant sequencing reads for SNVs, indels and deletions were visualised for validation on Integrative Genomes Viewer (IGV)^18^, and were confirmed by diagnostic capture-based high-throughput sequencing, if the IGV inspection was not satisfactory. For the familial segregation analysis, linkage to the *BMPR2* locus was first examined by microsatellite genotyping analysis. Mutation screening of the *BMPR2*, *ACVRL1*, *ENG*, *AQP1* and *SOX17* genes was conducted by capillary sequencing using BigDye Terminator v3.1 chemistry. All DNA fragments were resolved on an ABI Fragment Analyzer (Applied Biosystems). The family trees were drawn using the R package FamAgg^63^.

### Structural analysis of novel variants

The domain structures and the functional groups of the novel PAH genes were plotted according to the entry in UniProtKB. Clustal Omega was used for sequence alignment. Structural data were obtained from RCSB Protein Data Bank and analysed according to published reports. Figures were generated using PyMOL Molecular Graphics System.

### Production of pGDF2 Wild Type and Variant Proteins

The cloning of human wild type pro-GDF2 (pGDF2) in pCEP4 has been described previously^64^. Site-directed mutagenesis was performed according to the manufacturer’s instructions (QuickChange Site Directed Mutagenesis Kit, Agilent Technologies). Mutations were confirmed by Sanger sequencing. HEK-EBNA cells were transfected with plasmids containing either wild-type or mutant pGDF2 for 14 hours. The transfecting supernatant was removed and replaced with CDCHO media (Invitrogen) for 5 days to express the proteins. The conditioned media containing GDF2 and the variants were harvested and snap-frozen on dry-ice before being stored at −80^o^C. For each variant, conditioned media from three independent transfections were collected for further characterisation.

### GDF2 ELISA

High binding 96-well ELISA plates (Greiner, South Lanarkshire, UK) were coated with 0.2μg/well of mouse monoclonal anti-human GDF2 antibody (R&D Systems, Oxfordshire, UK) in PBS (0.1M phosphate pH7.4, 0.137M NaCl, 2.7mM KCl, Sigma) overnight at 4^o^C in a humidified chamber. Plates were washed with PBS containing 0.05% (v/v) Tween-20 (PBS-T), followed by blocking with 1% bovine serum albumin in PBS-T (1% BSA/PBS-T) for 90min at room temperature. Recombinant human GDF2 standards (1-3000pg/ml) or conditioned media samples (100μl/well of 1:30, 1:100, 1:300, 1:1000, 1:3000 and 1:10000 dilutions) were then added and incubated for 2h at room temperature. After washing, plates were then incubated with 0.04μg/well biotinylated goat anti-human GDF2 (R&D Systems) in 1% BSA/PBS-T for 2hr. Plates were washed, then incubated with ExtrAvidin(r)-Alkaline phosphatase (Sigma) diluted 1:400 in 1% BSA/PBS-T for 90 min. Plates were washed with PBS-T followed by water. The ELISA was developed with a colorimetric substrate comprising 1mg/ml 4-Nitrophenyl phosphate disodium salt hexahydrate (Sigma) in 1M Diethanolamine, pH9.8 containing 0.5mM MgCl_2_. The assay was developed in the dark at room temperature and the absorbance measured at 405nm.

### Cell culture and treatments

Distal human pulmonary artery smooth muscle cells (PASMCs) were cultured from explants of small pulmonary vessels (<2mm diameter) dissected from lung resection specimens. The lung parenchyma was separated from a pulmonary arteriole following the arteriolar tree to isolate 0.5-to 2-mm-diameter vessels, which were excised, cut into small fragments, plated in T25 flasks and left to adhere for 2 h. Once adhered, the explants were propagated in DMEM/20% FBS plus A/A until cells had grown out and were forming confluent monolayers. PASMCs were trypsinized, and subsequent passages were propagated in DMEM supplemented with 10% heat-inactivated FBS and A/A and maintained at 37^o^C in 95% air-5% CO2. The smooth muscle phenotype was confirmed by positive immunofluorescent staining using an antibody to smooth muscle specific alpha-actin (Clone IA4 Sigma-Aldrich; 1:100 dilution). A section of the pulmonary arteriole was collected, fixed in formalin and embedded in paraffin, and sections analysed to ensure that the vessel was of pulmonary origin. Papworth Hospital ethical review committee approved the use of the human tissues (Ethics Ref 08/H0304/56+5) and informed consent was obtained from all subjects.

Human blood outgrowth endothelial cells (BOECs) were generated from 40-80 ml peripheral blood isolated with informed consent from healthy controls. The study was approved by the Cambridgeshire 3 Research Ethics Committee, reference number 11/EE/0297. BOECs were cultured in EGM-2MV with 10% standard grade FBS (Life Technologies, Carlsbad, CA) and were used for experiments between passages 4 and 8^65^. The endothelial nature of BOECs was assessed by flow cytometry for endothelial surface marker expression as described previously^31^. All cell lines were routinely tested for mycoplasma contamination. For experiments involving BOEC generation,

Human pulmonary artery endothelial cells (PAECs) were purchased from Lonza (Cat. No. CC-2530; Basel, Switzerland) and maintained in EGM-2 with 2% fetal bovine serum (FBS; Lonza). PAECs were used between passages 4 and 8. For experimental studies, cells were treated with EBM-2 containing Antibiotic-Antimycotic (A/A; 100U ml^-1^ penicillin, 100 mg ml^-1^ streptomycin and 0.25 mg ml^-1^ amphotericin B, Invitrogen, Renfrewshire, UK) and FBS concentrations as stated. Cell lines were routinely tested for mycoplasma contamination and only used if negative.

### RNA preparation and quantitative reverse transcription-PCR

Total RNA was extracted using the RNeasy Mini Kit with DNAse digestion (Qiagen, West Sussex, UK). cDNA was prepared from 1 μg of RNA using the High Capacity Reverse Transcriptase kit (Applied Biosystems, Foster City, CA), according to the manufacturer’s instructions. All quantitative PCR reactions were prepared in MicroAmp optical 96-well reaction plates (Applied Biosystems) using 50 ng pl^−1^ cDNA with SYBR Green Jumpstart Taq Readymix (Sigma-Aldrich), ROX reference dye (Invitrogen) and custom sense and anti-sense primers (all 200 nmol l-^1^). Primers for human *ACTB* (encoding (3-actin), AQP1, ATP13A3, B2M, HPRT and SOX17 were designed using PrimerBLAST (https://www.ncbi.nlm.nih.gov/tools/primer-blast/) (Supplementary Table 12). Reactions were amplified on a Quantstudio 6 Real-Time PCR system (Applied Biosystems). The relative abundance of each target gene in different cell lines was compared using the equation 2-(CtG^OI^-Ct3HK) where Ct3HK corresponded to the arithmetic mean of the Cts for ACTB, B2M and HPRT for each sample. For expression analysis of siRNA knockdown, the 2-^(AACt)^ method was used and fold expression determined relative to the DH1 control.

### siRNA transfection

Prior to transfection, cells were preincubated in Opti-MEM-I reduced serum media (Invitrogen) for 2h before transfection with 10nM siRNA that had been lipoplexed for 20 min at RT with DharmaFECT1 (GE Dharmacon, Lafayette, CO). Cells were then incubated with the siRNA/DharmaFECT1 complexes for 4h at 37oC before replaced by full growth media. Cells were kept in growth media for 24h before further treatment. Knockdown efficiency was confirmed by mRNA expression or immunoblotting. For proliferation assays, parallel RNA samples were collected both on day0 and day6, confirming that *ATP13A3* expression was reduced by >90% on Day 0 and still reduced by >70% at Day 6. For all other assays, parallel RNA samples were collected on the day of the experiment to confirm knockdown, which was >90%. The siRNAs used were oligos targeting *ATP13A3* (SASI_Hs02_00356805) from Sigma-Aldrich and ON-TARGETplus non-targeting Pool (siCP; GE Dharmacon).

### Flow cytometric apoptosis assay

BOECs were plated 150,000/well into 6-well plates and transfected with si*ATP13A3* or siCP lipoplexed with DharmaFECT1. Cells were then serum-starved in EBM-2 (Lonza) containing 0.1% FBS and A/A for 8 hours before treating with EBM-2 and A/A containing either 0.1%FBS or 5%FBS for another 24 hours. Cells were then trypsinized and after washing with PBS, stained using the FITC Annexin V Apoptosis Dectection Kit I (BD Biosciences). For each condition, dual-staining of 5μl FITC conjugated Annexin V and 5μl propidium iodide (PI) were added and incubated at room temperature for 15 minutes. For the single staining controls for compensation, either 5μl FITC Annexin V or 5μl PI was added into non-transfected cells. All samples were analysed on BD Accuri^™^ C6 Plus platform (BD Biosciences). Data were collected and analysed using FlowJo software, with AnnexinV^+^/PI^-^cells defined as early apoptotic (Treestar).

### Caspase-Glo 3/7 assay

BOECs were seeded at a density of 150,000/well into 6-well plates and transfected with si*ATP13A3* or siCP lipoplexed with DharmaFECT1. For each condition, cells were trypsinized from 6-well plates and reseeded in triplicates into a 96-well plate at a density of 15,000-20,000/well and left to adhere overnight. Cells were quiesced in EBM-2 containing 0.1%FBS for 24h before treating with or without EBM-2 and A/A containing either 0.1%FBS or 5%FBS for 16 hours. For measuring caspase activities, 100ul Caspase-Glo^®^ 3/7 Reagent (G8091 Promega) was added into each well, incubated and mixed on a plate shaker in the dark for 30 minutes at room temperature. The lysates were transferred to a white-walled 96-well plate and luminescence was read in a GloMax^®^ luminometer (Promega).

### Accession codes

Data of PAH cases included in this manuscript and eligible for public release according to the UK Research Ethics rules have been deposited in the European Genome-phenome Archive (EGA) at the EMBL - European Bioinformatics Institute under accession number EGAD00001003423.

## Acknowledgments

We gratefully acknowledge the participation of patients recruited to the UK National Institute of Health Research BioResource - Rare Diseases (NIHR BR-RD) Study. We thank the NIHR BR-RD staff and co-ordination teams at the University of Cambridge, and the research nurses and coordinators at the specialist pulmonary hypertension centres involved in this study. The UK National Cohort of Idiopathic and Heritable PAH is supported by the NIHR BR-RD, the British Heart Foundation (BHF) (SP/12/12/29836), the BHF Cambridge Centre of Cardiovascular Research Excellence, the UK Medical Research Council (MR/K020919/1), the Dinosaur Trust, BHF Programme grants to RCT (RG/08/006/25302) and NWM (RG/13/4/30107), and the UK NIHR Cambridge Biomedical Research Centre. Funding for whole-exome sequencing was provided through a Bart’s Charity award (MGU0205) to RCT and DvH. NWM is a BHF Professor andNIHR Senior Investigator. CH is a NIHR Rare Disease Translational Research Collaboration Clinical PhD Fellow. LS is supported by the Wellcome Trust Institutional Strategic Support Fund (204809/Z/16/Z) awarded to St. George’s, University of London. CJR is supported by a BHF Intermediate Basic Science Research Fellowship (FS/15/59/31839). AL is supported by a BHF Senior Basic Science Research Fellowship (FS/13/48/30453). We acknowledge the support of the Imperial NIHR Clinical Research Facility, the Netherlands CardioVascular Research Initiative, the Dutch Heart Foundation, Dutch Federation of University Medical Centres, the Netherlands Organisation for Health Research and Development and the Royal Netherlands Academy of Sciences. We also gratefully acknowledge Dr Claudia Cabrera in the NIHR Barts Cardiovascular Biomedical Research Centre for bioinformatics support. We thank all the patients and their families who contributed to this research and the Pulmonary Hypertension Association (UK) for their support.

## Author contributions

S.G., N.W.M. and W.H.O. conceived and designed the research. S.G., M.H, M.B. and C.H. processed the data and performed the statistical analysis. S.G., M.H, M.B., C.H. and N.W.M. drafted the manuscript. L.S., R.D.M. and R.C.T. conducted the *SOX17* familial segregation analyses. W.L. performed the structural analysis of the novel causative variants. R.S. generated the mutant cells. J.H., R.M.S, B.L. and P.D.U. conducted the functional experiments on the novel disease genes. M.S. performed the immunohistochemistry for novel gene products. L.C.D. helped with the assessment of pertinent findings. O.S. was involved with data analysis. D.W. participated in DNA extraction, sample QC and plating. L.S, R.D.M, S.H, M.A., C.J.R., W.H.O., N.S., A.L., R.C.T. and M.R.W. helped with data analysis and interpretation and made critical revision of the manuscript for important intellectual content. J.M.M., C.M.T. and K.Y. coordinated data collection. N.W.M. and W.H.O. handled the funding for the study. All other authors were responsible for data acquisition and recruitment of subjects to the study and helped to draft the final version of the manuscript.

## References

1. Wagenvoort CA. The pathology of primary pulmonary hypertension. J Pathol. 1970 Aug;101(4):Pi

2. McGoon MD, Benza RL, Escribano-Subias P, Jiang X, Miller DP, Peacock AJ, et al. Pulmonary arterial hypertension: epidemiology and registries. J Am Coll Cardiol. 2013 Dec 24;62(25 Suppl):D51–9

3. McLaughlin VV, Shillington A, Rich S. Survival in primary pulmonary hypertension: the impact of epoprostenol therapy. Circulation. 2002 Sep 17;106(12):1477–82

4. Lane KB, Machado RD, Pauciulo MW, Thomson JR, Phillips JA 3rd, Loyd JE, et al. Heterozygous germline mutations in BMPR2, encoding a TGF-beta receptor, cause familial primary pulmonary hypertension. Nat Genet. 2000 Sep; 26(1):81–4

5. Deng Z, Morse JH, Slager SL, Cuervo N, Moore KJ, Venetos G, et al. Familial primary pulmonary hypertension (gene PPH1) is caused by mutations in the bone morphogenetic protein receptor-II gene. Am J Hum Genet. 2000 Sep;67(3):737–44

6. Evans JD, Girerd B, Montani D, Wang XJ, Galie N, Austin ED, et al. BMPR2 mutations and survival in pulmonary arterial hypertension: an individual participant data meta-analysis. Lancet Respir Med. 2016 Feb;4(2):129–37

7. Trembath RC, Thomson JR, Machado RD, Morgan NV, Atkinson C, Winship I, et al. Clinical and molecular genetic features of pulmonary hypertension in patients with hereditary hemorrhagic telangiectasia. N Engl J Med. 2001 Aug 2;345(5):325–34

8. Harrison RE, Berger R, Haworth SG, Tulloh R, Mache CJ, Morrell NW, et al. Transforming growth factor-beta receptor mutations and pulmonary arterial hypertension in childhood. Circulation. 2005 Feb 1;111(4):435–41

9. Nasim MT, Ogo T, Ahmed M, Randall R, Chowdhury HM, Snape KM, et al. Molecular genetic characterization of SMAD signaling molecules in pulmonary arterial hypertension. Hum Mutat. 2011 Dec;32(12):1385–9

10. Shintani M, Yagi H, Nakayama T, Saji T, Matsuoka R. A new nonsense mutation of SMAD8 associated with pulmonary arterial hypertension. J Med Genet. 2009 May;46(5):331–7

11. Austin ED, Ma L, LeDuc C, Berman Rosenzweig E, Borczuk A, Phillips JA 3rd, et al. Whole exome sequencing to identify a novel gene (caveolin-1) associated with human pulmonary arterial hypertension. Circ Cardiovasc Genet. 2012 Jun;5(3):336–43

12. Ma L, Roman-Campos D, Austin ED, Eyries M, Sampson KS, Soubrier F, et al. A novel channelopathy in pulmonary arterial hypertension. N Engl J Med. 2013 Jul 25;369(4):351–361

13. Kerstjens-Frederikse WS, Bongers EM, Roofthooft MT, Leter EM, Douwes JM, Van Dijk A, et al. TBX4 mutations (small patella syndrome) are associated with childhood-onset pulmonary arterial hypertension. J Med Genet. 2013 Aug;50(8):500–6

14. Eyries M, Montani D, Girerd B, Perret C, Leroy A, Lonjou C, et al. EIF2AK4 mutations cause pulmonary veno-occlusive disease, a recessive form of pulmonary hypertension. Nat Genet. 2014 Jan;46(1):65–9

15. Best DH, Sumner KL, Austin ED, Chung WK, Brown LM, Borczuk AC, et al. EIF2AK4 mutations in pulmonary capillary hemangiomatosis. Chest. 2014 Feb;145(2):231–236

16. Abenhaim L, Moride Y, Brenot F, Rich S, Benichou J, Kurz X, et al. Appetite-suppressant drugs and the risk of primary pulmonary hypertension. International Primary Pulmonary Hypertension Study Group. N Engl J Med. 1996 Aug 29;335(9):609–16

17. Machado RD, Southgate L, Eichstaedt CA, Aldred MA, Austin ED, Best DH, et al. Pulmonary Arterial Hypertension: A Current Perspective on Established and Emerging Molecular Genetic Defects. Hum Mutat. 2015 Dec;36(12):1113–27

18. Thorvaldsdottir H, Robinson JT, Mesirov JP. Integrative Genomics Viewer (IGV): high-performance genomics data visualization and exploration. Brief Bioinform. 2013 Mar;14(2):178–92

19. Hadinnapola C, Bleda M, Haimel M, Screaton N, Swift A, Dorfmuller P, et al. Phenotypic Characterization of EIF2AK4 Mutation Carriers in a Large Cohort of Patients Diagnosed Clinically With Pulmonary Arterial Hypertension. Circulation. 2017 Nov 21;136(21):2022–2033

20. Hamid R, Cogan JD, Hedges LK, Austin E, Phillips JA 3rd, Newman JH, et al. Penetrance of pulmonary arterial hypertension is modulated by the expression of normal BMPR2 allele. Hum Mutat. 2009 Apr;30(4):649–54

21. David L, Mallet C, Mazerbourg S, Feige JJ, Bailly S. Identification of BMP9 and BMP10 as functional activators of the orphan activin receptor-like kinase 1 (ALK1) in endothelial cells. Blood. 2007 Mar 1;109(5):1953–61

22. Mi LZ, Brown CT, Gao Y, Tian Y, Le VQ, Walz T, et al. Structure of bone morphogenetic protein 9 procomplex. Proc Natl Acad Sci U S A. 2015 Mar 24;112(12):3710–5

23. David L, Mallet C, Keramidas M, Lamande N, Gasc JM, Dupuis-Girod S, et al. Bone morphogenetic protein-9 is a circulating vascular quiescence factor. Circ Res. 2008 Apr 25;102(8):914–22

24. Thever MD, Saier MH Jr. Bioinformatic characterization of p-type ATPases encoded within the fully sequenced genomes of 26 eukaryotes. J Membr Biol. 2009 Jun;229(3):115–30

25. Kanai R, Ogawa H, Vilsen B, Cornelius F, Toyoshima C. Crystal structure of a Na+-bound Na+,K+-ATPase preceding the E1P state. Nature. 2013 Oct 10;502(7470):201–6

26. Sui H, Han BG, Lee JK, Walian P, Jap BK. Structural basis of water-specific transport through the AQP1 water channel. Nature. 2001 Dec 20-27;414(6866):872–8

27. Sinner D, Rankin S, Lee M, Zorn AM. Sox17 and beta-catenin cooperate to regulate the transcription of endodermal genes. Development. 2004 Jul;131(13):3069–80

28. Remenyi A, Lins K, Nissen LJ, Reinbold R, Scholer HR, Wilmanns M. Crystal structure of a POU/HMG/DNA ternary complex suggests differential assembly of Oct4 and Sox2 on two enhancers. Genes Dev. 2003 Aug 15;17(16):2048–59

29. Palasingam P, Jauch R, Ng CK, Kolatkar PR. The structure of Sox17 bound to DNA reveals a conserved bending topology but selective protein interaction platforms. J Mol Biol. 2009 May 8;388(3):619–30

30. Jauch R, Aksoy I, Hutchins AP, Ng CK, Tian XF, Chen J, et al. Conversion of Sox17 into a pluripotency reprogramming factor by reengineering its association with Oct4 on DNA. Stem Cells. 2011 Jun;29(6):940–51

31. Toshner M, Dunmore BJ, McKinney EF, Southwood M, Caruso P, Upton PD, et al. Transcript analysis reveals a specific HOX signature associated with positional identity of human endothelial cells. PLoS One. 2014;9(3):e91334

32. Schultheis PJ, Hagen TT, O’Toole KK, Tachibana A, Burke CR, McGill DL, et al. Characterization of the P5 subfamily of P-type transport ATPases in mice. Biochem Biophys Res Commun. 2004 Oct 22;323(3):731–8

33. Madan M, Patel A, Skruber K, Geerts D, Altomare DA, Iv OP. ATP13A3 and caveolin-1 as potential biomarkers for difluoromethylornithine-based therapies in pancreatic cancers. Am J Cancer Res. 2016;6(6):1231–52

34. Taraseviciene-Stewart L, Kasahara Y, Alger L, Hirth P, Mc Mahon G, Waltenberger J, et al. Inhibition of the VEGF receptor 2 combined with chronic hypoxia causes cell death-dependent pulmonary endothelial cell proliferation and severe pulmonary hypertension. FASEB J. 2001 Feb;15(2):427–38

35. Teichert-Kuliszewska K, Kutryk MJ, Kuliszewski MA, Karoubi G, Courtman DW, Zucco L, et al. Bone morphogenetic protein receptor-2 signaling promotes pulmonary arterial endothelial cell survival: implications for loss-of-function mutations in the pathogenesis of pulmonary hypertension. Circ Res. 2006 Feb 3;98(2):209–17

36. Wang G, Fan R, Ji R, Zou W, Penny DJ, Varghese NP, et al. Novel homozygous BMP9 nonsense mutation causes pulmonary arterial hypertension: a case report. BMC Pulm Med. 2016 Jan 22;16:17

37. Long L, Ormiston ML, Yang X, Southwood M, Graf S, Machado RD, et al. Selective enhancement of endothelial BMPR-II with BMP9 reverses pulmonary arterial hypertension. Nat Med. 2015 Jul;21(7):777–85

38. Saadoun S, Papadopoulos MC, Hara-Chikuma M, Verkman AS. Impairment of angiogenesis and cell migration by targeted aquaporin-1 gene disruption. Nature. 2005 Apr 7;434(7034):786–92

39. Schuoler C, Haider TJ, Leuenberger C, Vogel J, Ostergaard L, Kwapiszewska G, et al. Aquaporin 1 controls the functional phenotype of pulmonary smooth muscle cells in hypoxia-induced pulmonary hypertension. Basic Res Cardiol. 2017 May;112(3):30

40. Matsui T, Kanai-Azuma M, Hara K, Matoba S, Hiramatsu R, Kawakami H, et al. Redundant roles of Sox17 and Sox18 in postnatal angiogenesis in mice. J Cell Sci. 2006 Sep 1;119(Pt 17):3513–26

41. Corada M, Orsenigo F, Morini MF, Pitulescu ME, Bhat G, Nyqvist D, et al. Sox17 is indispensable for acquisition and maintenance of arterial identity. Nat Commun. 2013;4:2609

42. Lange AW, Haitchi HM, LeCras TD, Sridharan A, Xu Y, Wert SE, et al. Sox17 is required for normal pulmonary vascular morphogenesis. Dev Biol. 2014 Mar 1;387(1):109–20

43. Sundin OH, Leppert GS, Silva ED, Yang JM, Dharmaraj S, Maumenee IH, et al. Extreme hyperopia is the result of null mutations in MFRP, which encodes a Frizzled-related protein. Proc Natl Acad Sci U S A. 2005 Jul 5;102(27):9553–8

44. The Genotype-Tissue Expression (GTEx) project. Nat Genet. 2013 Jun;45(6):580–5

45. Machado RD, Pauciulo MW, Thomson JR, Lane KB, Morgan NV, Wheeler L, et al. BMPR2 haploinsufficiency as the inherited molecular mechanism for primary pulmonary hypertension. Am J Hum Genet. 2001 Jan;68(1):92–102

46. DePristo MA, Banks E, Poplin R, Garimella KV, Maguire JR, Hartl C, et al. A framework for variation discovery and genotyping using next-generation DNA sequencing data. Nat Genet. 2011 May;43(5):491–8

47. Yang H, Wang K. Genomic variant annotation and prioritization with ANNOVAR and wANNOVAR. Nat Protoc. 2015 Oct;10(10):1556–66

48. Lek M, Karczewski KJ, Minikel EV, Samocha KE, Banks E, Fennell T, et al. Analysis of protein-coding genetic variation in 60,706 humans. Nature. 2016 Aug 18;536(7616):285–91

49. Auton A, Brooks LD, Durbin RM, Garrison EP, Kang HM, Korbel JO, et al. A global reference for human genetic variation. Nature. 2015 Oct 1;526(7571):68–74

50. Walter K, Min JL, Huang J, Crooks L, Memari Y, McCarthy S, et al. The UK10K project identifies rare variants in health and disease. Nature. 2015 Oct 1;526(7571):82–90

51. Kircher M, Witten DM, Jain P, O’Roak BJ, Cooper GM, Shendure J. A general framework for estimating the relative pathogenicity of human genetic variants. Nat Genet. 2014 Mar;46(3):310–5

52. Ng PC, Henikoff S. Predicting deleterious amino acid substitutions. Genome Res. 2001 May;11(5):863–74

53. Adzhubei IA, Schmidt S, Peshkin L, Ramensky VE, Gerasimova A, Bork P, et al. A method and server for predicting damaging missense mutations. Nat Methods. 2010 Apr;7(4):248–9

54. Cooper GM, Stone EA, Asimenos G, Green ED, Batzoglou S, Sidow A. Distribution and intensity of constraint in mammalian genomic sequence. Genome Res. 2005 Jul;15(7):901–13

55. Conomos MP, Miller MB, Thornton TA. Robust inference of population structure for ancestry prediction and correction of stratification in the presence of relatedness. Genet Epidemiol. 2015 May;39(4):276–93

56. Conomos MP, Reiner AP, Weir BS, Thornton TA. Model-free Estimation of Recent Genetic Relatedness. Am J Hum Genet. 2016 Jan 7;98(1):127–48

57. Purcell S, Neale B, Todd-Brown K, Thomas L, Ferreira MA, Bender D, et al. PLINK: a tool set for whole-genome association and population-based linkage analyses. Am J Hum Genet. 2007 Sep;81(3):559–75

58. Li H, Handsaker B, Wysoker A, Fennell T, Ruan J, Homer N, et al. The Sequence Alignment/Map format and SAMtools. Bioinformatics. 2009 Aug 15;25(16):2078–9

59. Zhan X, Hu Y, Li B, Abecasis GR, Liu DJ. RVTESTS: an efficient and comprehensive tool for rare variant association analysis using sequence data. Bioinformatics. 2016 May 1;32(9):1423–6

60. Chen X, Schulz-Trieglaff O, Shaw R, Barnes B, Schlesinger F, Kallberg M, et al. Manta: rapid detection of structural variants and indels for germline and cancer sequencing applications. Bioinformatics. 2016 Apr 15;32(8):1220–2

61. Roller E, Ivakhno S, Lee S, Royce T, Tanner S. Canvas: versatile and scalable detection of copy number variants. Bioinformatics. 2016 Aug 1;32(15):2375–7

62. Zarrei M, MacDonald JR, Merico D, Scherer SW. A copy number variation map of the human genome. Nat Rev Genet. 2015 Mar;16(3):172–83

63. Rainer J, Taliun D, D’Elia Y, Pattaro C, Domingues FS, Weichenberger CX. FamAgg: an R package to evaluate familial aggregation of traits in large pedigrees. Bioinformatics. 2016 May 15;32(10):1583–5

64. Wei Z, Salmon RM, Upton PD, Morrell NW, Li W. Regulation of bone morphogenetic protein 9 (BMP9) by redox-dependent proteolysis. J Biol Chem. 2014 Nov 7;289(45):31150–9

65. Ormiston ML, Toshner MR, Kiskin FN, Huang CJ, Groves E, Morrell NW, et al. Generation and Culture of Blood Outgrowth Endothelial Cells from Human Peripheral Blood. J Vis Exp. 2015 Dec 23;(106):e53384

